# Targetron-assisted delivery of exogenous DNA sequences into *Pseudomonas putida* through CRISPR-aided counterselection

**DOI:** 10.1101/2021.05.01.442236

**Authors:** Elena Velázquez, Yamal Al-Ramahi, Jonathan Tellechea, Natalio Krasnogor, Víctor de Lorenzo

## Abstract

Genome editing methods based on Group II introns (known as Targetron technology) have been long used as a gene knock-out strategy in a wide range of organisms in a fashion independent of homologous recombination. Yet, their utility as delivery systems has been typically suboptimal because of their reduced efficiency of insertion when they carry exogenous sequences. We show that this limitation can be tackled and Targetron adapted as a general tool in Gram-negative bacteria. To this end, a set of broad host range standardized vectors were designed for conditional expression of the Ll.LtrB intron. After testing the correct functionality of these plasmids in *Escherichia coli* and *Pseudomonas putida*, we created a library of Ll.LtrB variants carrying cargo DNA sequences of different lengths to benchmark the capacity of intron-mediated delivery in these bacteria. Next, we combined CRISPR/Cas9-facilitated counterselection to increase the chances of finding genomic sites inserted with the thereby engineered introns. By following this pipeline, we were able to insert exogenous sequences of up to 600 bp at specific genomic locations in wild-type *P. putida* KT2440 and its Δ*recA* derivative. Finally, we were able to apply this technology to successfully tag this strain with an orthogonal short sequence (barcode) that acts as a unique identifier for tracking this microorganism in biotechnological settings. The results with *P. putida* exemplified the value of the Targetron approach for unrestricted delivery of small DNA fragments to the genomes of Gram-negative bacteria for a suite of genome editing endeavours.

---

*Pseudomonas putida* is a soil bacterium and plant root colonizer that has emerged as one of the species with the highest potential as a Synthetic Biology chassis for industrial and environmental applications^1,2^. Qualities of interest include the lack of pathogenicity ^3^, its high tolerance to oxidative stress^4,5^ (a most desirable trait in processes such as biofuel production^6^), diverse and powerful capabilities for catabolizing aromatic compounds^7–9^ and ease of genetic and genomic manipulations^10–13^. In particular, a suite of molecular tools have become available for deletion and insertion of foreign sequences in the genome of this soil bacterium, both randomly (e.g. with transposon vectors^14,15^) and directed to specific genomic loci through recombineering^16^ or homologous recombination^13^ (reviewed in^17^). In this last and most widely used case, note that recombination efficacies vary considerably among different bacterial groups and even strains of the same species—the archetypal *P. putida* KT2440 specimen being one particularly suboptimal in *recA*-dependent processes.

Group II introns could be a way to overcome this problem. These molecules are a type of retroelements with the capacity to splice from an mRNA and insert into specific DNA loci (a process known as retrohoming^18,19^. Their conserved structure and a protein codified by themselves (IEP or Intron-Encoded Protein) are key components for the splicing and recognition of the target DNA^20,21^. After translation, the IEP binds specifically to the intronic RNA and assists its splicing process from the exons. Afterwards, both molecules keep attached forming a ribonucleoprotein (RNP) that will carry out the recognition, reverse splicing as well as retrotranscription of the intronic RNA into the new DNA molecule^22^. In the past, several of these introns have been engineered to recognize and insert into specific genes different from their native retrohoming sites, giving rise to the knock-out system named Targetron^23^.

Targetron is founded on Ll.LtrB group II intron from *Lactococcus lactis* since it is the most studied intron of this class and was proven to work in a wide range of bacterial genres, from *Clostridium*^24^ or *Bacillus*^25^ to the well-characterized species *Escherichia coli*^23^. Later, Targetron was also surveyed to be exploited as a delivery system of cargos into designated loci^26–28^. Nevertheless, this attempt highlighted the most serious limitations group II introns have. First, they can be modified to recognize new sequences but their integration efficiency can greatly change depending on the new target site. Indeed, mathematical algorithms have been developed to identify the best retargeting options in a given sequence for Ll.LtrB^23,24^ and also for other group II introns^29^. These algorithms retrieve a list of loci ordered by a predicted score and they also design primers for the modification of the recognition sequences inside the intron. However, as a result of their probabilistic nature, these predictions are not always reliable. Secondly, cargo sequences can be inserted inside of group II introns to be transported. In fact, the optimal region inside of the intron to carry these cargos has been greatly studied. it was described how domain IVb was the best insertion point and Ll.LtrB was modified to display a MluI restriction site at this position^27^. Yet, the presence of exogenous sequences in this domain also hinders the efficiency of group II to splice and retrohome to some extent. Therefore, despite the good characteristics of these molecules, their possibility to be boosted as delivery systems was poorly achieved. Recently, some efforts to overcome these drawbacks were made when CRISPR/Cas9 technology was merged with Targetron to ease the identification of invaded mutants in *E. coli*^30^. This was accomplished by directing Cas9 endonuclease to the insertion site of Ll.LtrB with the help of specific spacers recognizing this area. Thereby, if the intron retrohomed into the correct locus, CRISPR/Cas9 will no longer couple with this region and the invaded mutant will survive. On the other hand, if the intron did not insert, Cas9 will cleave the bacterial genome and these cells will die. However, the applicability of these two systems in other species has not been deeply addressed as well as the total capacity of this combination to increase the size of fragments that can be delivered. In this context, the generation of a sensitive, broad-host expression system compatible with CRISPR/Cas9 plasmids is required.

In the work described below, we have generated a set of SEVA plasmids expressing Ll.LtrB intron under the control of different promoters (IPTG and cyclohexanone induction^31^), origins of replication as well as antibiotic resistance genes that work in a broad-host-range of Gram-negative bacteria. To test the correct behaviour of the new expression plasmids, we have engineered them to insert Ll.LtrB intron into different genes of *E. coli* and *P. putida*. Next, we cloned a library of sequences with different sizes inside pSEVA6511-GIIi to address the possibility of coupling the CRISPR/Cas9 system ^32^ as a counterselection mechanism for Ll.LtrB insertions in *P. putida* KT2440. Besides, we have selected this Gram-negative soil bacterium and its *recA* mutant counterpart to validate the utility of this system in strains with little or non-existent homologous recombination^33^. Finally, we used this technology to successfully label *P. putida* KT2440 with a specific synthetic barcode that could identify and trace down this strain in future applications^34^. The data presented here not only shows the functionality of the generated system but also its behaviour in a new microorganism in which Targetron technology had not been assayed before.

## RESULTS AND DISCUSSION

### Engineering broad-host expression of Ll.LtrB intron

Ll.LtrB intron has been exploited to work in diverse organisms from bacteria or yeast to mammalian cells by using different expression systems. Commercial Targetron technology has been validated in a wide range of bacterial species such as *Staphylococcus aureus*^35^, *E. coli*^23^, *L. lactis*^27^, *Shigella flexneri*^36^, *Salmonella typhimurium*^36^ or *Clostridium perfringens*^24^. However, even with this proven broad-host functionality of Ll.LtrB, Targetron has the drawback of having to re-clone and adapt the backbone to the organism at stake that is to be engineered. This is why there have been several attempts to build broad-host-range plasmids that could work in general Gram-negative bacteria. For instance, a mini-RK2 plasmid with tetracycline resistance was engineered to express Ll.LtrB from the XylS/*Pm* promoter and its activity was surveyed in species such as *E. coli, P. aeruginosa* and *Agrobacterium tumefaciens*^37^.

The SEVA (Standard European Vector Architecture)^12,38,39^ database was launched as one attempt in Synthetic Biology to establish a standardized and coherent collection of plasmids with both a minimalist format and nomenclature. The first set of these formatted vectors is composed of different interchangeable modules including broad-host range origins of replication, antibiotic resistance genes and a wide set of expression systems and reporter genes. In this context, we decided to couple the standardization and robustness of SEVA plasmids with the Targetron technology. To this end, we engineered a collection of pSEVAs to express Ll.LtrB intron so that different induction strategies could be chosen freely according to the sought purpose. First, pSEVA421-GIIi (Km) (Supplementary Fig. S1) was generated and tested in *E. coli* BL21DE3 (Fig. 1A and 1B). This plasmid has a low copy number (RK2 origin of replication), streptomycin/spectinomycin resistance and the Ll.LtrB intron and LtrA (Ll.LtrB IEP) sequences under control of a T7 promoter. The cloned intron was retargeted to insert into the *lacZ* gene so that blue/white screening could be used to assess the accuracy of the insertion process. In addition to this, Ll.LtrB in this plasmid also carries a retrotransposition-activated selectable marker (RAM) in domain IVb which has been previously reviewed to increase the likelihood of finding retrohomed mutants^40^. RAM in pSEVA421-GIIi(Km) is composed of Km^R^ gene interrupted by a group I intron. The construct is arranged in a way that only if the intron is inserted, the group I intron is excised and the Km^R^ gene is reconstituted. Therefore, selection in Km plates facilitates the identification of insertion mutants (Fig. 1A, right plates).

**Figure 1.**
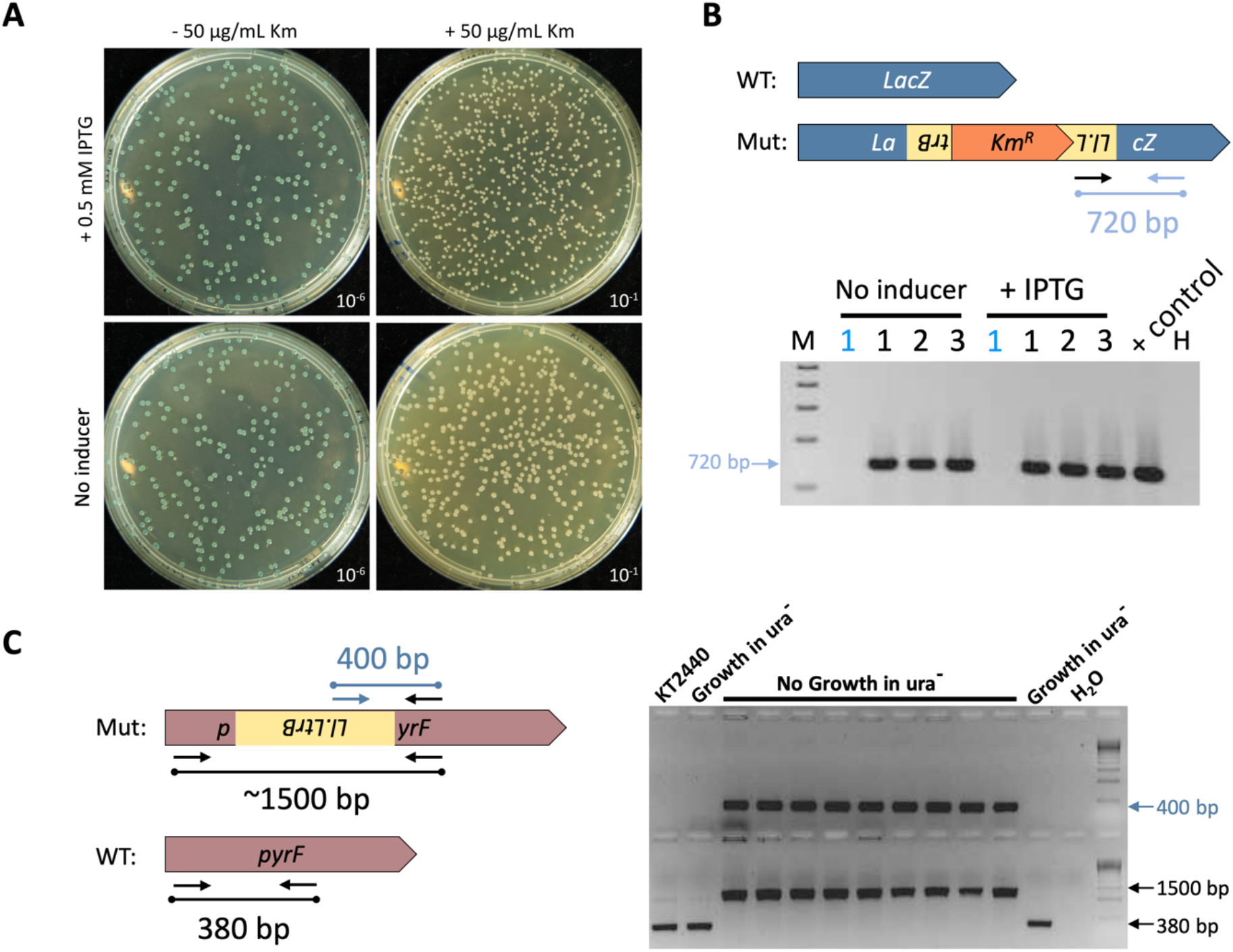
SEVA plasmids encoding Ll.LtrB group II intron and T7RNAP work in A,B) *E. coli* BL21•DE3 and C) *P. putida* KT2440. A) Delivery of Ll.LtrB intron from plasmid pSEVA421-GIIi(Km) in *E. coli* BL21DE3. Ll.LtrB intron was retargeted to insert into locus 1063a of *lacZ* gene so that insertions would disrupt this gene, giving rise to white colonies in the presence of X-gal. Since a RAM is placed inside Ll.LtrB, kanamycin resistance could be used as a way to select intron insertion mutants (plates to the right). B) PCR reactions to determine the correct insertion of Ll.LtrB inside *lacZ* gene. Only if Ll.LtrB retrohomes, a PCR amplicon of 720bp is generated. Blue numbers correspond to blue colonies and black numbers correspond to white colonies used as the template material for each reaction. C) Delivery of Ll.LtrB from plasmid pSEVA421-GIIi-pyrF and with help of pSEVA131-T7RNAP in *P. putida* KT2440. 5FOA counterselection was used to isolate insertion mutants that were not able to grow without uracil supplemented to plates. Colonies resistant to 5FOA but that were able to grow without uracil were used as negative controls of insertion. Two different PCR reactions are shown: (Top gel) one primer annealed inside Ll.LtrB and the second annealed in the *pyrF* gene so that an amplicon could be only generated after intron insertion. (Bottom gel) two primers flanking the insertion locus were used so that two amplicons could be generated. The smallest fragment (380bp) corresponds to the WT sequence and the biggest fragment (1500bp) corresponds to the insertion. The same colonies were tested in both PCR reactions. WT: Wild-type, Mut: Insertion Mutant, + control: reaction with an invaded colony from a previous experiment used as template, H_2_O: Control PCR with no template material.

Since a T7 promoter controls the expression of Ll.LtrB, T7 RNA polymerase (T7RNAP) is needed for transcription. This is why we first checked the activity of this plasmid in *E. coli* BL21DE3 as it has a bacteriophage *λ* derivative (prophage DE3) with the T7RNAP gene under the control of the lacUV5 promoter^41^. The results of the blue/white screening in the insertion assay performed with pSEVA421-GIIi (Km) showed that the intron seemed to be retrohoming to the selected locus inside the *lacZ* gene (Fig. 1A). It also highlights the importance of having method to spot insertion mutants. When Km was supplemented to plates, the number of white colonies was undoubtedly boosted in comparison to the plates with no selection (Fig. 1A, left plates). Moreover, we observed a slight increase in the number of white colonies when adding the inducer IPTG (~1.5 fold). Finally, colony PCR of white and blue colonies was performed to check the correct insertion of the intron in the *lacZ* gene and the correlation with the disclosed phenotype (Fig. 1B).

### Ll.LtrB intron retrohomes in *P. putida* KT2440

After testing the efficacy of the new SEVA plasmids, we decided to check the activity of Ll.LtrB in *P. putida* since no previous work has studied the performance of this group II intron in this species. First, we needed to engineer a new pSEVA for the heterologous expression of the T7RNAP. The complete sequence of this ORF, along with the regulatory regions for IPTG-controlled expression (lacUV5 promoter along with a short 5’ region of *lacZ* gene fused to the T7RNAP ORF), were cloned into pSEVA131, yielding pSEVA131-T7RNAP (Supplementary Fig. S1). This plasmid bears an ampicillin resistance gene (Ap^R^), a pBBR1 origin of replication which confers a medium copy number of plasmids and, most importantly, is compatible with pSEVA421-GIIi(Km). On the other hand, a *lacZ* gene ortholog is not present in the genome of *P. putida* KT2440. Therefore, we needed to search for a different reporter gene of insertion. A very useful genetic marker that has been widely employed in positive and negative selection is the gene URA3 and its homologs. URA3 encodes the orotidine-5’-phosphate decarboxylase (ODCase) which is an enzyme that participates in the biosynthetic pathway of pyrimidines in *Saccharomyces cerevisiae*^42^. Thereby, inactivation of this gene leads to uracil auxotrophy that can be complemented by adding this pyrimidine in media. On the other hand, ODCase also catalyzes the transformation of 5-fluoroorotic acid (5FOA) into 5-fluorouracil, a toxic compound that causes cell death^43^. Therefore, negative selection (loss of ODCase activity) is based on the growth of URA3-disrupted mutants in the presence of 5FOA and uracil in media. Instead, positive selection works based on complementing the loss of ODCase with an active gene that can be supplemented through a plasmid or any other exogenous construct. Given that *P. putida* KT2440 is endowed with an ortholog of URA3 called *pyrF* (PP1815) and that this gene has been already used several times as a counterselection marker in this strain^44^, we decided that this gene would be an ideal candidate for Ll.LtrB insertion.

Once the target gene was chosen, Ll.LtrB was retargeted to insert into one specific locus inside this ORF, giving rise to pSEVA421-GIIi(Km)-pyrF. A variant of this plasmid, pSEVA421-GIIi-pyrF, was also generated. The only difference was the absence of RAM since 5FOA-based counterselection could be applied instead of directly selecting for successfully retrohomed mutants based on Km resistance. Both plasmids were transformed into *P. putida* KT2440, respectively, along with pSEVA131-T7RNAP and the insertion assay was performed as in *E. coli*. The only exception was that a longer incubation time (2 h) was chosen to ensure enough expression levels from both plasmids. After selection in plates supplemented with 5FOA and uracil, growing cells were patched on plates with and without uracil to search for real *pyrF* mutants. In the case of pSEVA421-GIIi-pyrF, colonies growing on 5FOA/uracil and being real uracil auxotrophs were identified. Two PCR reactions were set to verify the presence of Ll.LtrB in the *pyrF* gene at the correct site (Fig. 1C). This result demonstrated the ability of Ll.LtrB to retrohome inside *P. putida* KT2440 as it has been shown in other species of this genre like *P. aeruginosa* ^37^. Interestingly, we were not able to identify insertion events when using Ll.LtrB::RAM by using either Km or 5FOA selection. Even though the use of RAMs has been validated in different organisms like *E. coli* ^40^ and *L. lactis* ^45^, the impossibility to find insertion mutants with this selection method was also reported to happen in *P. aeruginosa*^37^. In that work, the authors stated the possibility of this to be caused by the lack of processivity of RNA polymerases from hosts. That is, RNA polymerases might not be able to transcribe the whole sequence of group II introns containing cargos as they would disclose a long and complex structure. Nevertheless, the authors also envisioned the option to overcome this limitation by supplying T7RNAP whose processivity and transcription frequency has been previously demonstrated^41,46,47^. Nevertheless, in our experimental setup, we used this approach and no mutants were found. Besides, the activity of the pSEVA131-T7RNAP was demonstrated after finding Ll.LtrB insertions by using 5FOA counterselection. This leads us to a second possibility that can be related to either the excision of the group I intron present inside the RAM or other problems related to the relative efficiency of splicing of the intron in this species.

### Simplification of the Ll.LtrB expression system and its activity in the Δ*recA* derivative strain of *P. putida* KT2440

Even if both engineered pSEVAs are functional in general Gram-negative bacteria, having to transform two plasmids inside the strain to be modified remains a hindrance. With this in mind, we decided to try out a new and simpler expression system that could alleviate this necessity. By sub-cloning Ll.LtrB and LtrA from pSEVA421-GIIi-pyrF into pSEVA2311^31^, we generated pSEVA2311-GIIi-pyrF (Supplementary Fig. S1). With a Km^R^ and a pBBR1 *oriV*, this vector controls the expression of Ll.LtrB from a ChnR/P_ChnB_ promoter regulated by the addition of the aromatic compound cyclohexanone. This backbone has been previously validated in *E. coli*^31^ and also employed to regulate biofilm formation in *P. putida* ^48^. We expected sustained Ll.LtrB expression in Gram-negative bacteria under this promoter.

The same 5FOA-mediated counterselection was used to isolate insertion mutants in both wild-type *P. putida* KT2440 and its Δ*recA* derivative (Fig. 2). Different concentrations of cyclohexanone were employed (0, 0.5, 1 and 5 mM) and, in all cases, we were able to retrieve clones with the retrohomed intron in both strains. This supports the ability of Ll.LtrB intron to work in a recombinant-independent fashion in *P. putida*. This is a feature of group II introns that had been already observed previously in other species ^33^,^49^.

**Figure 2.**
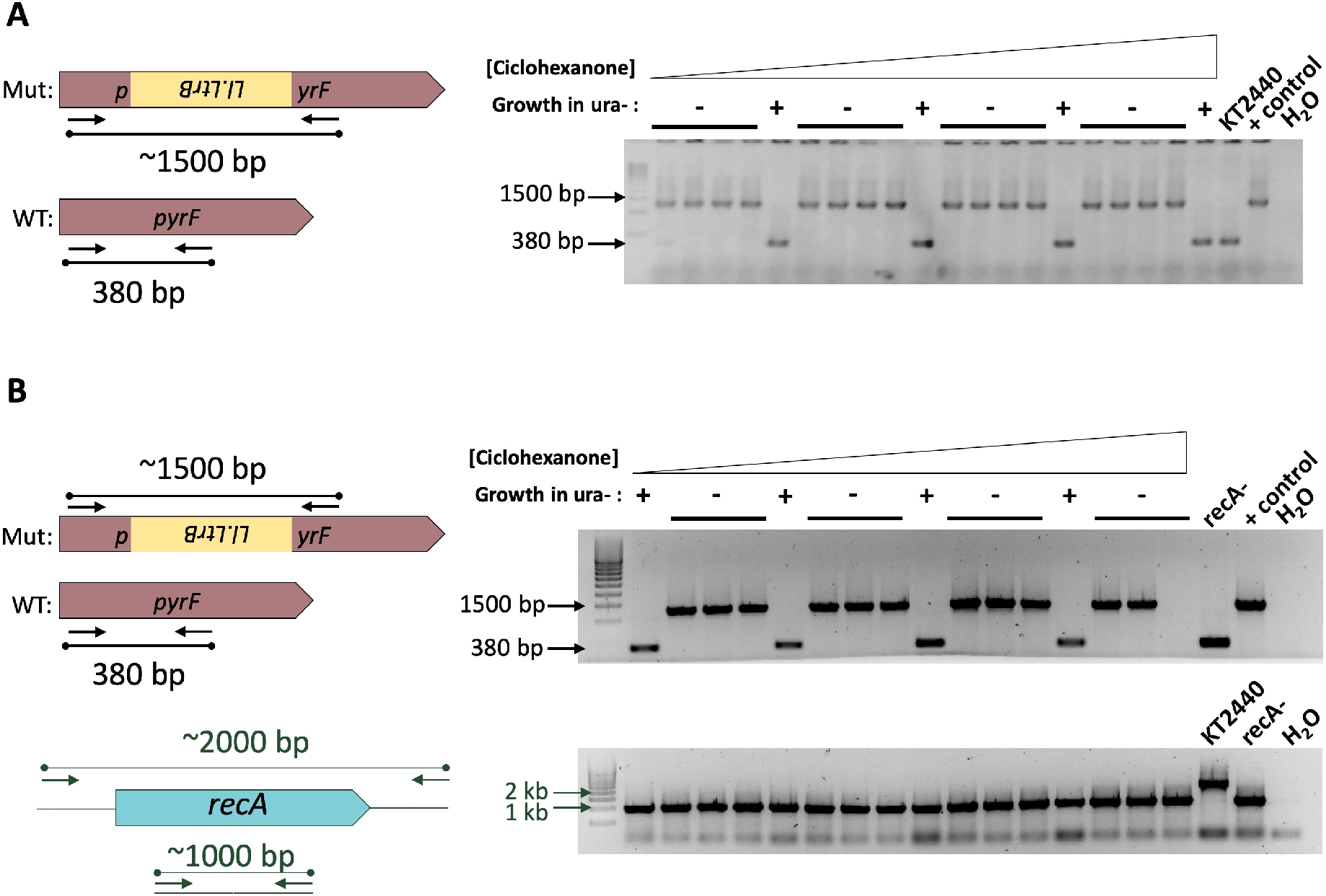
Performance of pSEVA2311-GIIi in *P. putida* KT2440 and its Δ*recA* derivative strain. A) pSEVA2311-GIIi works in *P. putida* KT2440 WT to deliver Ll.LtrB intron into *pyrF* gene through 5FOA counterselection. Different concentrations of cyclohexanone were used from 0mM (left part of gel) to 5mM (right part of gel). A PCR reaction that used primers flanking the insertion locus was used to determine Ll.LtrB retrohoming. The smallest fragment (380bp) corresponds to the WT sequence while the biggest one (1500bp) corresponds to the intron insertion. B) pSEVA2311-GII also works in *P. putida* KT2440 Δ*recA*. (Top gel) the same PCR reaction with flanking primers was performed to determine intron insertions. No amplification was considered as a negative result for Ll.LtrB insertion and 5FOA resistance was considered to be due to another mechanisms which could be affecting the amplification during PCR (i.e. possible deletion of part/entire *pyrF* gene). (Bottom gel) PCR reaction to verify *recA* minus genotype of the tested cells. The same colonies were tested in both PCR reactions. WT: Wild-type, Mut: Insertion Mutant, KT2440: Parental *P. putida* KT2440 WT was used as template material, recA^-^: Parental *P. putida* KT2440 Δ*recA* was used as template material, + control: reaction with an invaded colony from a previous experiment used as template, H_2_O: Control PCR with no template material.

### Construction of a high copy number plasmid expressing Ll.LtrB compatible with CRISPR/Cas9-mediated counterselection system

Our next goal was generating a new pSEVA with both oriV and antibiotic marker compatible with the CRISPR/Cas9-mediated counterselection engineered previously in our laboratory ^30,32^. For this, we built pSEVA6511-GIIi and pSEVA6511-GIIi(Km) (Supplementary Fig. S1). Both plasmids have a high copy number origin of replication (RSF1010), a gentamycin resistance gene and a *lacZ*-retargeted Ll.LtrB and LtrA controlled by the ChnR/p_ChnB_ promoter.

The efficacy of these plasmids to deliver Ll.LtrB was again tested in *E. coli* BL21DE3 (Fig. 3A) and *P. putida* KT2440 (Fig. 3B), in this case, after being retargeted towards *pyrF* gene. In the *E. coli* strain, we were able to identify insertion mutants in both cases, i.e., with and without Km selection. On the contrary, in the case of *P. putida*, we were still unable to find retrohomed mutants when using Km^R^ RAM. Still, empty Ll.LtrB was able to invade *pyrF* which proved the functioning of pSEVA6511-GIIi-pyrF in this microorganism. This last experiment also highlighted the level of spontaneous mutations arising to 5FOA that have already been characterized ^44^ and the necessity of streaking 5FOA-resistant colonies to verify their uracil auxotrophy and thus the disruption of the *pyrF* gene. Even so, with no striking, a moderate frequency of mutants was detected just with 5FOA selection (3 colonies out of 16, which gives a frequency ~ 18 %). Accordingly, after testing colonies that were unable to grow without uracil in media, the frequency of detected insertion mutant went up to ~ 75% (9 out of 12 colonies).

**Figure 3.**
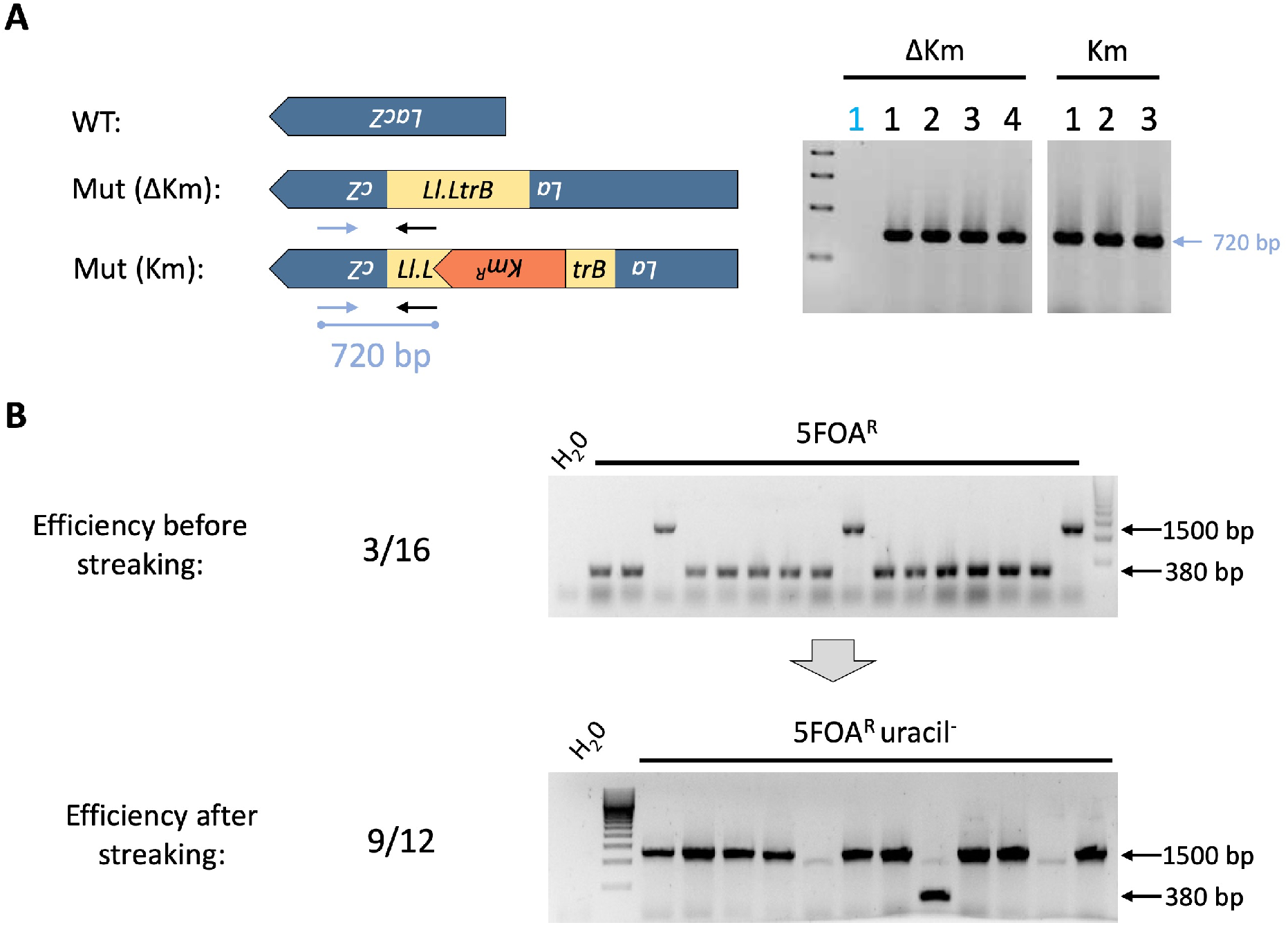
Engineered pSEVA6511-GIIi expresses correctly Ll.LtrB intron in both A) *E. coli* BL21DE3 and B) *P. putida* KT2440. A) pSEVA6511-GIIi(Ø/Km) plasmids express Ll.LtrB intron in *E. coli* BL21DE3 and inserts correctly inside *lacZ* gene. A PCR reaction where a primer anneals inside Ll.LtrB and the other anneals in the *lacZ* gene was employed in both cases to verify the correct insertion of the intron. A fragment of 720bp is generated when Ll.LtrB is present in the selected locus. Blue numbers correspond to blue colonies and black numbers correspond to white colonies used as the template material for each reaction B) pSEVA6511-GIIi-pyrF works in *P. putida* to deliver Ll.LtrB intron into the *pyrF* gene with 5FOA counterselection (top gel) and uracil auxotrophy (bottom gel). The same PCR reaction using primers flanking the insertion locus inside *pyrF* gene were used. An amplicon of 380bp is generated if Ll.LtrB is not present (WT) while a fragment of 1500bp is amplified if the intron is present (Mut). WT: Wild-type, Mut: Insertion Mutant, H_2_O: Control PCR with no template material.

### Limits in the size of fragments that can be delivered by Ll.LtrB intron in *P. putida* KT2440 and its Δ*recA* derivative

Once pSEVA6511-GIIi-pyrF was checked to work in *P. putida*, we proceeded to survey the idea of exploiting Ll.LtrB for exogenous sequences delivery into specific loci in the genome of this soil bacterium. Group II introns have been previously studied with this approach in mind in different organisms ^27,28,30,50^. Different types of cargos have been employed in these works as well as different target integration sites. In general, all of them concluded that the size of the sequences inserted inside Ll.LtrB was critical for the splicing and retrohoming efficiency of the intron. In fact, in its native host, *L. lactis*, cargo sequences longer than 1 kb started to highly hinder the frequency of detected insertion mutants^27^. Between the results showed above, we stated the unfeasibility of finding Ll.LtrB::RAM mutants in *P. putida*. Since the size of this RAM was 1.2 kb, this corresponds with the observations of previous works where this fragment length causes Ll.LtrB insertions to be undetectable in some cases. Nevertheless, this same Ll.LtrB::RAM was interestingly able to efficiently retrohomed in *E. coli* (Fig. 1A and 3A) which made us think that this limitation in size could be host-dependent. For this reason, we decided to study the maximum length of a fragment that Ll.LtrB was able to carry in both WT and ΔrecA *P. putida* KT2440. To do so, we generated a library of pSEVA6511-GIIi-pyrF plasmids carrying fragments of increasing size from the luxC gene (Fig. 4A). After this, we perform the same insertion assay with and without using 5FOA-mediated counterselection with insert sizes of 150 bp (Lux1), 600 bp (Lux4), 750 bp (Lux5) and 1050 (Lux7). In the case of WT *P. putida* KT2440, we were able to identify insertion mutants with Lux4 when using 5FOA counterselection. Nonetheless, insertions with Lux5 or Lux7 were not detected (Supplementary Fig. S2, and Supplementary Table S3). In the case of not using any type of counterselection, we could only identify one insertion mutant out of 100 colonies in the case of Lux1, making clear the utility of having a counterselection mechanism. *P. putida* Δ*recA* only showed sign of insertion with the smallest size tested (Supplementary Table S3) which was something surprising as no differences were expected between the two strains regarding Ll.LtrB intron mobility.

**Figure 4.**
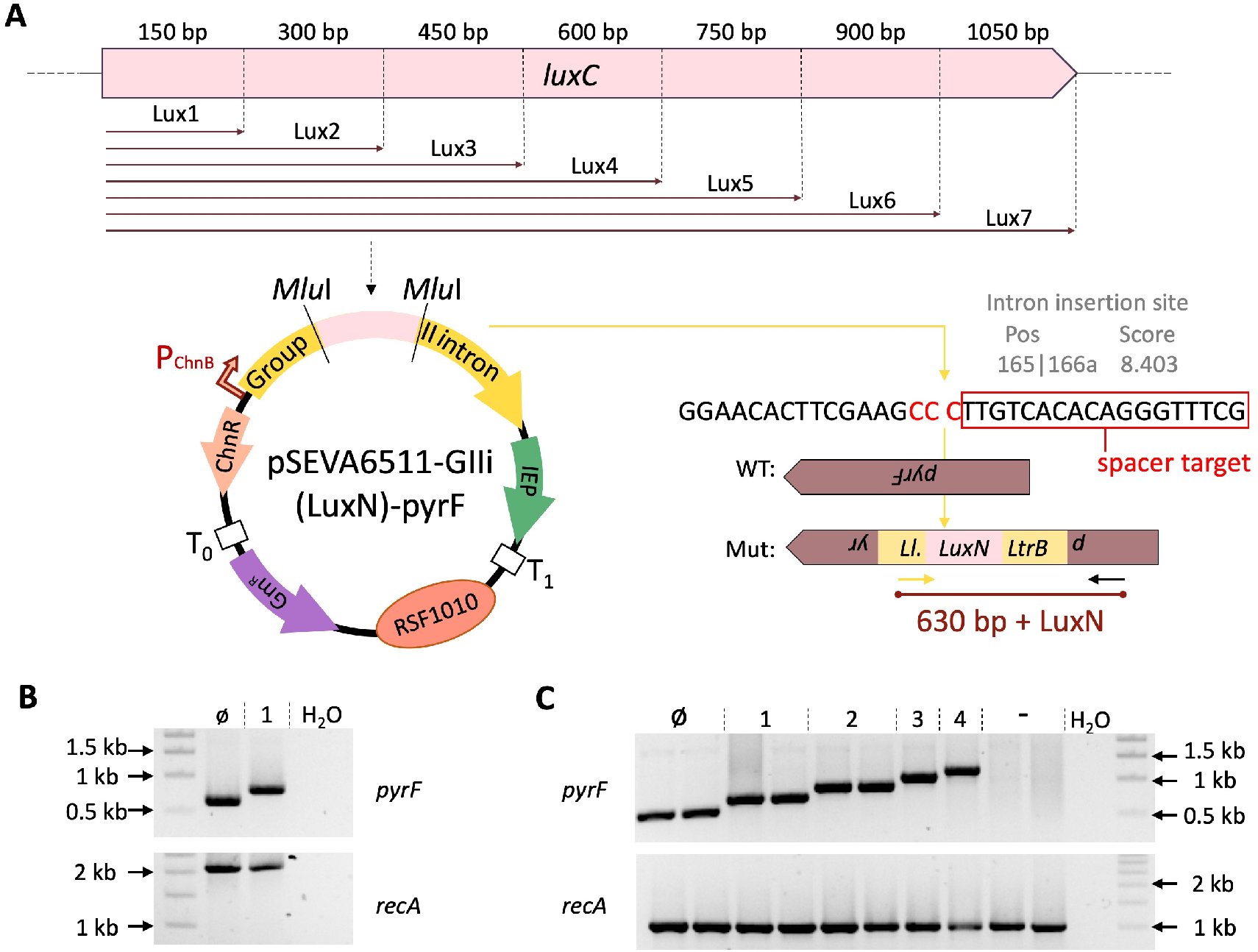
Assessing size-restriction of intron-mediated delivery using *luxC* fragments as cargo A) Schematic of the intron library generated with increasing fragment length as cargo (from 150bp up to 1050bp) using as template the first gene of the luxCDABEG operon, *luxC*. Ll.LtrB intron in pSEVA6511-GIIi(LuxN) is retargeted to insert between the nucleotides 165 and 166 of the *pyrF* ORF in the antisense orientation. Spacer pyrF1 recognizes the region after the insertion site (part of the recognition site is shown inside a red box). The complementary nucleotides to the PAM (5’ -GGG −3’) are highlighted in red. B) L.LtrB-mediated delivery of *luxC* fragments in *P. putida* KT2440 WT. PCR reaction showing amplifications from colonies with Ll.LtrB::LuxØ and Ll.LtrB::Lux1 (top gel) and PCR reaction showing the *recA* genotype (bottom gel). WT amplification for *recA* gen is 2 kb long. C) Ll.LtrB-mediated delivery of *luxC* fragments in *P. putida* KT2440 ΔrecA. PCR reaction showing amplifications from colonies with Ll.LtrB::LuxØ to Ll.LtrB::Lux4 (top gel) and PCR reaction showing the *recA* genotype (bottom gel). Deletion of the *recA* gene gives an amplification of 1 kb. WT: Wild-type, Mut: Insertion Mutant, LuxN: Cargos including from LuxØ to Lux7, Ø: Ll.LtrB with no cargo, 1: Ll.LtrB with Lux1 as cargo,2: Ll.LtrB with Lux2 as cargo; 3: Ll.LtrB with Lux3 as cargo, 4: Ll.LtrB with Lux4 as cargo, -: *P. putida* KT2440 Δ*recA* colonies with no inserted Ll.LtrB used as a negative control.

### Incorporating CRISPR/Cas9-mediated counterselection of Ll.LtrB intron insertion

CRISPR/Cas9-based counterselection has proven to act as a perfect tool to increase the efficiency of different mutagenesis procedure by eliminating the WT population of non-modified cells^32,51,52^. CRISPR machinery has been described as an adaptive immune system in bacteria ^53^ and it is composed of two main elements: the CRISPR array and the <u>C</u>RISPR-<u>ass</u>ociated (Cas) proteins. The first component comprises the <u>C</u>lustered <u>R</u>egularly <u>I</u>nterspaced <u>S</u>hort <u>P</u>alindromic <u>R</u>epeats (CRISPR itself) and specific spacers that are placed between these repeats. Regarding Cas proteins, the most outstanding one is called Cas9, a double-stranded DNA endonuclease that has been greatly employed in CRISPR-mediated counterselection ^32,51^. As the general mechanism, when these two components are expressed, the spacers are complexed with Cas9 and guide the endonuclease to specific regions in the genome by base pairing. If their recognition target is closed to a <u>P</u>rostospacer-<u>A</u>dajacent <u>M</u>otive (PAM), Cas9 will cleave that genomic locus. In this context, by designing spacers that can couple with WT sequences and cleave them to cause cell death, the likelihood to identify specific mutants in the selected target was improved, and, as a result, a counterselection method was generated.

In previous work, we were successfully able to couple this counterselection mechanism with Targetron technology in *E. coli* ^30^. By designing specific spacers that recognize the WT sequence at the insertion site of group II introns, we were able to facilitate the identification of invaded mutants. As the next step in our study, we tried to apply this same procedure in *P. putida*. The objective was to test if this counterselection mechanism could help us to boost the size limit of the fragments that could be delivered with Ll.LtrB. Cells harbouring both pSEVA421-Cas9tr ^32^ and the corresponding pSEVA6511-GIIi(LuxN)-pyrF were grown overnight and then induced for 4h with cyclohexanone. After this incubation time, an aliquot of these cells was plated in the presence of 5FOA to estimate the efficiency of intron insertions with this induction protocol and no CRISPR counterselection. The rest of the cells were made competent and then, two conditions were tested: Either pSEVA231-CRISPR (a negative control with no specific spacer) or pSEVA231-C-pyrF1 (with a specific spacer recognizing the insertion locus of Ll.LtrB::LuxN) were transformed into respective aliquots. The last plasmid bears a spacer that has been already assayed for *pyrF*-mutants counterselection in *P. putida*^32^. If Ll.LtrB::LuxN retrohomes, the PAM sequence necessary for Cas9 activation will be disrupted and mutated cells will be able to survive (Fig. 4A).

By following this approach, we were able to identify mutated cells in both *P. putida* WT (Fig. 4B and Fig. 5A, B) and Δr*ecA* (Fig. 4C and Fig. 5C, D). Nevertheless, we could still not find insertions for fragments longer than 600 bp (Lux4). In addition to this, this time we were able to isolate mutants with this fragment size in the Δ*recA* background (Fig. 4C) while we could only identify insertions coming from Ll.LtrB::LuxØ and Ll.LtrB::Lux1 in *P. putida* WT (Fig. 4B). Regarding the insertion frequencies comparing all conditions, the general amount of detected integrations was incremented when transforming pSEVA231-C-pyrF1 in both backgrounds (Fig. 5 A, C and Supplementary Tables S4 and S5). Besides, with no CRISPR spacer, no insertions with Ll.LtrB::Lux2-4 were spotted after surveying a total of ~100 colonies per condition when transforming pSEVA231-CRISPR (Supplementary Table S5). Taken together, these results indicate that the system allows the positive selection of the LL.LtrB integrations by counterselecting the population of non-mutated bacteria. The low increase in this efficiency in comparison with the results obtained previously with *E. coli* ^30^ could be explained by the differences in the targets selected in each case. In this matter, changing the insertion locus and CRISPR spacer could help to further increase the number of mutants and even help to identify integrations with longer fragments.

**Figure 5.**
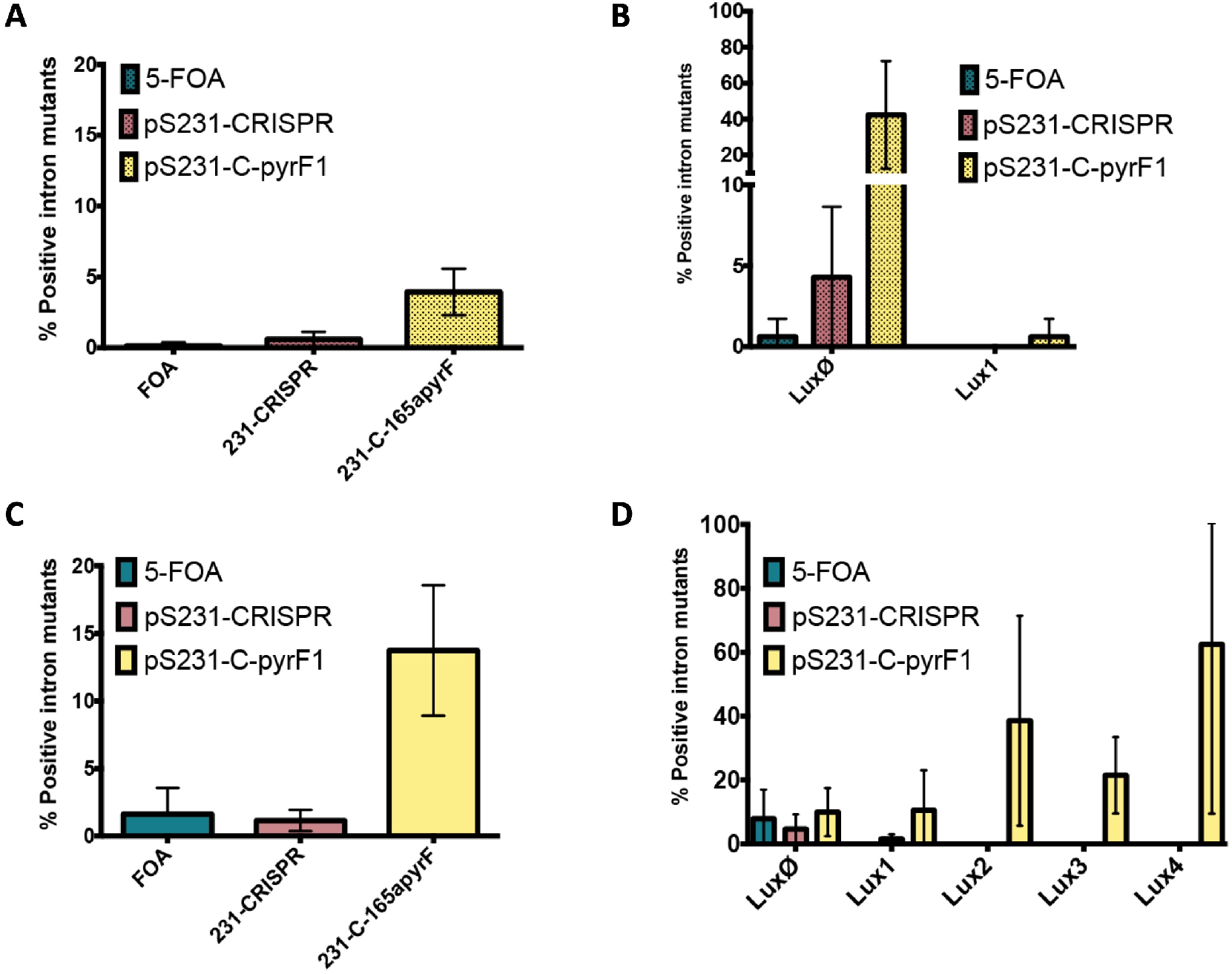
Intron insertion frequencies in *P. putida* KT2440 WT and ΔrecA confirmed through PCR. A) Total intron-insertion frequency in *P. putida* KT2440 WT using 5FOA or CRISPR/Cas9-mediated counterselection. B) Intron-insertion frequency of each cargo being delivered to the genome of *P. putida* KT2440 WT by Ll.LtrB using 5FOA or CRISPR/Cas9-mediated counterselection. C) Total intron-insertion frequency in *P. putida* KT2440 ΔrecA using 5FOA or CRISPR/Cas9-mediated counterselection. D) Intron-insertion frequency of each cargo being delivered to the genome of *P. putida* KT2440 ΔrecA by Ll.LtrB using 5FOA or CRISPR/Cas9-mediated counterselection. The average and standard deviation of two or three replicates are shown. Ø: Ll.LtrB with no cargo; 1: Ll.LtrB with Lux1 as cargo,2: Ll.LtrB with Lux2 as cargo; 3: Ll.LtrB with Lux3 as cargo, 4: Ll.LtrB with Lux4 as cargo.

### Application of Ll.LtrB for the delivery of small genetic fragments that can be used as traceable barcodes

Tracing genetically modified strains along with their pedigrees and modifications can be very challenging. Nevertheless, this is a critical step for the sake of archiving and also for biosafety in case we happen to deliver those strains into the environment. One solution for this could be the use of small pieces of synthetic DNA (a.k.a. genetic barcodes) that could serve as identifiers of particular strains. By introducing these barcodes in the genomes of modified cells, we can create a physical link between this engineered organism and its digital counterpart. This idea has been described previously along with a version control system for microbial strains (named CellRepo) where all important information about barcoded strains can be archived and consulted in repositories^34,54^. In this way, after barcoding one strain, one only needs to sequence the barcode to retrieve all the available information about that strain on the website, including laboratory of origin, developer, modifications, resistances, etc.

As we have shown above along with data previously published about the performance of Ll.LtrB intron in other organisms^27^, group II introns seems to be particularly useful for the delivery of small fragments of DNA. Therefore, we thought they could be an optimum tool for the delivery of orthogonal sequences as barcodes/unique identifiers to the genomes of desired strains as they have a small size (148 bp, Supplementary Fig. S3). As proposed in^34^, an optimal barcode structure is composed by a universal primer (25 nt), which is shared by all barcodes generated with the CellRepo software, and a core sequence (123 nt) which is subdivided into three components: the barcode sequence itself (96 nt), the synchronization (9 nt) and the checksum (18 nt) sequences^34^. The last two elements are incorporated as an error-correction mechanism. This way, even if truncated or incorrect reads are retrieved, the CellRepo algorithm is still able to identify the barcode and its linked strain profile content. On the other hand, some of the main features of group II introns are that, first, they insert themselves stably in DNA molecules as it has been proven for them to generate stable integrations after more than 80 generations^28,50^. Second, as it was already stated, they can function in a wide range of hosts^24,25,27,35,36^. Third, group II introns can be retargeted to insert into virtually any desired loci of choice and they have high specificity for their target ^23,29^. Finally, as we have also shown, they are independent of the RecA-based homologous recombination machinery, which is an advantage compared to other apprioaches to the same end ^30,33^. Considering all this, we decided to examine Ll.LtrB as the carrier of a specific barcode to label *P. putida* KT2440. First, we synthesized the specific barcode generated with the CellRepo algorithm by using two overlapping oligonucleotides (Supplementary Fig. S3) and then we cloned it into pSEVA6511-GIIi to generate pSEVA6511-GIIi(B3).

Next, we searched for good targeting loci in the genome of *P. putida*. As barcodes are meant to link a strain to its digital data, they need to be included in a stable and favourable genetic locus. This is why we chose intergenic regions close to well-known essential genes in *P. putida*. Thereby, *glmS* context was selected as a good candidate for the insertion of Ll.LtrB::B3. Two different insertion loci were selected from the retrieved list after using Clostron algorithm to survey the intergenic region between *PP5408* and *glmS* (Fig. 6 and Supplementary Fig. S4). Then, Ll.LtrB was retargeted towards these two loci, giving rise to pSEVA6511-GIIi(B3)-37s and pSEVA6511-GIIi(B3)-94a, respectively (Fig. 6). Next, specific spacers for both loci were designed and tested to see the efficiency of cleavage which was about one order of magnitude in both cases (Supplementary Fig. S5). After these two components were ready, we performed the same insertion protocol we had already established for the delivery of *luxC* fragments. Obtained colonies were directly checked through pool PCR reactions to analyze a high number of colonies at once as no phenotype change was expected after the insertion of Ll.LtrB. Only in the case of locus 2, insertion mutants were located with the corresponding barcode sequence (Fig. 7). Additional PCRs were performed to secure the purity of the final mutated colony (data not shown) and the barcode integrity was confirmed through sequencing (Details about the final barcoded strain can be found in the public CellRepo repository https://cellrepo.ico2s.org/repositories/93?branch_id=139&locale=en). This final result emphasizes the utility of having an external counterselection system to improve the screening of Ll.LtrB insertions as, in most cases, a phenotype change is not expected in thereby watermarked cells. Moreover, in this last case, the scores predicted for both loci (4.438 and 1.719, Supplementary Fig. S4) were much lower than those obtained for the *lacZ* (9.188) and *pyrF* (8.403) gene. It is been previously stated in the text how this score is just a prediction and cannot be blindly relied on. However, it is important to consider it to fully assess the utility of the CRISPR/counterselection system in these cases where a low efficiency of insertion is to be expected.

**Figure 6.**
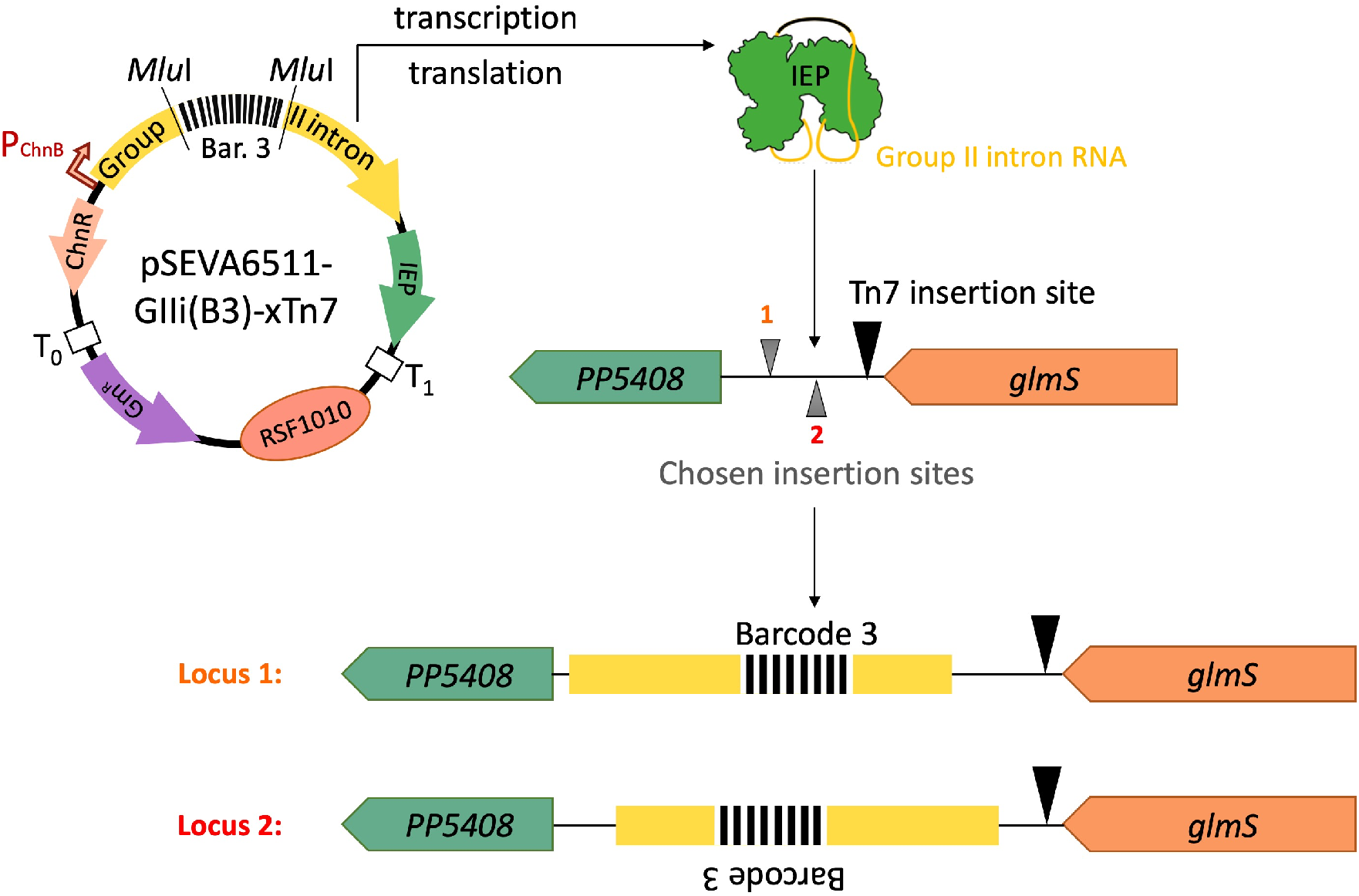
Application of Ll.LtrB group II intron for the delivery of specific genetic barcodes to the genome of *P. putida* KT2440 WT. Selection of the insertion loci for Ll.LtrB::B3 in the vicinity of the Tn7-insertion site (black triangle). Two different insertion points (grey triangles) were chosen for the insertion list generated in the Clostron website and Ll.LtrB::B3 was retargeted to both sites accordingly. The recognition site in Locus 1 (orange) is located in the sense strand while Locus 2 (red) is present in the antisense strand of *P. putida’*s genome. Ll.LtrB::B3 insertion would generate two different genotypes depending on the locus being targeted in each case.

**Figure 7.**
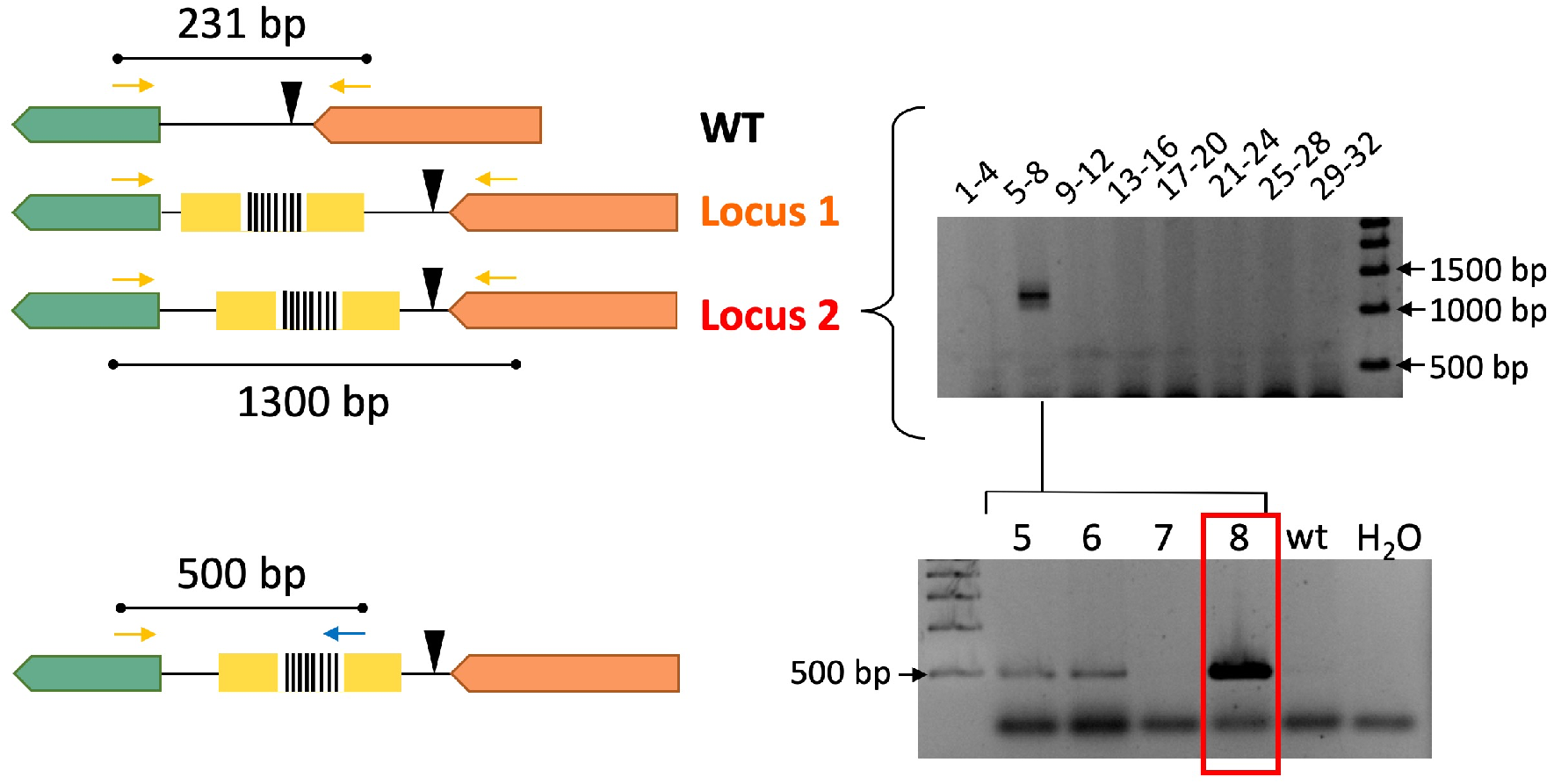
Delivery of Ll.LtrB::B3 into *P. putida* KT2440 WT genome. A first pool PCR was set to detect Ll.LtrB::B3 successful insertions in either Locus 1 (orange) or locus 2 (red). The top gel shows the amplification found with a pool PCR using primers flanking the insertion locus 2. The bottom gel shows the second PCR of individual colonies from the corresponding pool to find the barcoded clone. In this case, a primer annealing inside the barcode (pbarcode universal) and other annealing inside *PP5408* gene were used. WT: Wild-type, Locus 1: 37,37s insertion site, Locus 2: 94,95a insertion site. *PP5408* (green gene), *glmS* (orange gene).

### Conclusion

While *P. putida* KT2440 is a strain that has made evident its utility in biotechnological applications, there is still a need to develop new tools that can be used to modify this bacterium and broaden its applicability. Moreover, it is important to build these new tools in a broad-host-range manner that would allow their use in other bacteria with little modification. Our work tried to expand the number of genetic devices that can be exploited to insert sequences of certain length at specific genomic regions by using group II introns. Also, a wide-host format was adopted by expressing Ll.LtrB intron from different SEVA plasmids that are functional in general Gram-negative bacteria. The efficacy of these pSEVAs was tested in *E. coli* and *P. putida* KT2440 WT as well as in its derivative Δ*recA*, highlighting one of the main advantages of this technology which is its independence of homologous recombination. Finally, we also validated the possibility of coupling group II introns and CRISPR/Cas9-mediated counterselection in *P. putida* to improve the efficiency of searching for insertion mutants and demonstrated its use in the labelling of particular strains with genetic barcodes.

## METHODS

### Bacterial strains and media

*E. coli* CC118 strain [Δ(*ara-leu*), *araD139*, *ΔlacX74, galE, galK phoA20, thi-1, rpsE, rpoB, argE* (Am), *recA1*, OmpC^+^, OmpF^+^] was used for plasmid cloning and propagation and BL21DE3 strain [*fhuA2*, [lon], *ompT, gal*, (λ DE3), [dcm], Δ*hsdS*; (λ DE3) = λ sBamHIo ΔEcoRI-B int::(*lacI::PlacUV5*::T7 gene1) i21 Δnin5] for intron mobility assays in *E. coli. P. putida* KT2440 and its derivative Δ*recA* were used to assess intron mobility in this specie._Luria-Bertani (LB) medium was used for general growth and was supplemented when needed with kanamycin (Km; 50 μg/mL), ampicillin (Ap; 150 μg/mL for *E. coli* and 500 μg/mL for *P. putida*), gentamycin (Gm; 10 μg/mL for *E. coli* and 15 μg/mL for *P. putida*) and/or streptomycin (Sm; 50 μg/mL)._For solid plates, LB medium was supplemented with 1.5% of agar (w/v). In specific cases for *P. putida*, M9 minimal medium [6 g L^-1^ Na2HPO4, 3 g L^-1^ KH2PO4, 1.4 g L^-1^ (NH4)2SO4, 0.5 g L^-1^ NaCl, 0.2 g L^-1^ MgSO4·7H2O] supplemented with sodium citrate at 0.2% (w/v) as the carbon source was used instead. X-gal (5-Bromo-4-chloro-3-indolyl-β-D-galactopyranoside) was added at a final concentration of 30 μg/mL to carry out blue/white colony screening._Moreover, different inducers were added to media when necessary: isopropyl-1-thio-b-galactopyranoside (IPTG) was added at 0.5 mM (*E. coli*) or 1mM (*P. putida*) and cyclohexanone at 1 mM if not stated differently.

### Plasmid construction

The complete sequence encoding the T7 promoter, Ll.LtrB intron and LtrA protein was amplified from the commercial plasmid pACD4K-C (TargeTron gene knockout system, Sigma-Aldrich) with primers pGIIintron_fwd and rev (Supplementary Table S1). The amplified fragment was then digested with PacI and SpeI restriction enzymes and cloned into a similarly digested pSEVA427, yielding pSEVA421-GIIi(Km). On the other hand, *lacUV5* promoter along with T7 RNA polymerase (T7RNAP) sequences were amplified from pAR1219 (Merck, Sigma-Aldrich) with primers pAR1219_fwd and rev, PacI/SpeI digested and cloned into corresponding sites of pSEVA131, generating pSEVA131-T7RNAP, necessary for the transcription of Ll.LtrB intron from the T7 promoter. To eliminate the retrotransposition-activated selectable marker (RAM) present inside Ll.LtrB, pSEVA421-GIIi(Km) was digested with MluI restriction enzyme and then directly ligated and transformed to obtain pSEVA421-GIIi._To change the expression system and simplify the intron expression mechanism, only Ll.LtrB (with or without RAM) and LtrA sequences were extracted by HindIII/SpeI digestion of pSEVA421-GIIi and cloned into pSEVA2311, giving rise to pSEVA2311-GIIi(Km) and pSEVA2311-GIIi, respectively. These plasmids have both Ll.LtrB intron and LtrA expression controlled under the ChnR-P_ChnB_ promoter. Finally, to assemble an expression plasmid compatible with the CRISPR/Cas9 system described previously ^32^, it was necessary to modify both the origin of replication and the antibiotic resistance gene. For that, the ChnR-P_ChnB_ promoter, Ll.LtrB (with and without RAM) and LtrA sequences were extracted by digestion with PacI/SpeI enzymes and cloned into pSEVA651 equivalent sites to obtain pSEVA6511-GIIi(Km) and pSEVA6511-GIIi. The CRISPR/Cas9 counterselection approach used in this work was described in ^32^ and is based on plasmids pSEVA421-Cas9tr and pSEVA231-CRISPR. pSEVA231-C-pyrF1 was generated and described in the same work. The rest of the spacers for counterselection were designed manually and cloned into BsaI-digested pSEVA231-CRISPR, following the protocol explained in the same paper. The resulting plasmids were named pSEVA231-C-37s and pSEVA231-C-94a.

### Retargeting of Ll.LtrB intron

Retargeting of Ll.LtrB intron was performed by adapting the Targetron protocol from Sigma-Aldrich. First, Clostron platform was used to design primers pIBS-X, pEBS1d-X, pEBS2-X and pEBSuniversal (depending on the insertion target; Supplementary Table S1) with corresponding target sequences as query (*lacZ* gene in *E. coli; pyrF* gene and PP5408-glmS region in *P. putida*). From the output list, the best-ranked targets compatible with CRISPR/Cas9 technology were selected in each case. This means targets with PAM sequences (5’-NGG-3’ in the case of Streptococcus pyogenes system) closest to the insertion site of the intron were chosen. Afterwards, Clostron-designed oligonucleotides for each target were used in a SOEing PCR with pACD4K-C as a template to yield a 350bp fragment. For the cloning of this amplicon, different strategies were adopted attending to the final recipient plasmid. For the retargeting of pSEVA421-GIIi and its derivatives, the fragment was digested with BsrGI/HindIII restriction enzymes and ligated into the linearized recipient plasmid. For retargeting of pSEVA2311-GIIi and pSEVA6511 derivatives, Gibson assembly was chosen as the cloning procedure since an addition BsrGI restriction sites was present in the p_ChnB_ promoter. Primers pRetarget-fwd and rev were used to reamplify the SOEing amplicon and add the corresponding homologous sequences to directly assemble the fragment to HindIII/HpaI-digested pSEVA2311/6511-GIIi.

### Insertion of exogenous sequences inside Ll.LtrB

All exogenous sequences inserted inside Ll.LtrB intron were cloned into the MluI site present in the intron sequence. In the case of the insert to be delivered, two strategies were followed: *LuxC* gene from the lux operon was employed as a template for the generation of fragments of different sizes (from 150bp to 1050bp with a difference of 150bp each). Primer pLux_fwd in combination with primers pLux1-7_rev (Supplementary Table S1) respectively were used in a PCR step to generate each fragment using as template pSEVA256. Each amplicon was then digested with MluI and cloned into linearized pSEVA6511-GIIi-pyrF. The orientation of each fragment was confirmed by sequencing. Barcodes sequences were created in the CellRepo website (https://cellrepo.herokuapp.com) with an algorithm that provides universally unique identifiers (UUIDs) (ref Natalio). This provides the possibility to produce a large library of barcodes randomly generated and unique. After selecting one specific barcode, a BLAST search was done to make sure there was no other region with high similarity in the genome of *P. putida*. Once a barcode was verified, it was generated by a PCR step with 119-mer oligonucleotides bearing 30 overlapping nucleotides at 3’. These primers also included 30 nucleotides complementary to the recipient vector at 5’, so Gibson assembly reaction could be performed after amplification with MluI-digested pSEVA6511-GIIi.

### Interference assay of spacers 37s and 94a

*P. putida* KT2440 strain harbouring pSEVA421-Cas9tr was grown overnight and electrocompetent cells were prepared by washing cells with 300mM sucrose a total of 5 times. The final pellet was resuspended on 400 μL and then split into 100 μL aliquots. One hundred nanograms of pSEVA231-CRISPR (control), pSEVA231-C-37s or 94a (Supplementary Table S2) were electroporated into respective aliquots. Transformed bacteria were grown in LB/Sm for 2h at 30°C and serial dilutions were then plated on LB/Sm to test viability and LB/Sm/Km plates to assess the efficiency of cleavage. After counting CFUs on both conditions, the ratio of transformation efficiency was calculated by dividing the CFUs on LB/Sm plates by CFUs on LB/Sm/Km plates both normalized to 10^9^ cells.

### Ll.LtrB insertion assay in *E. coli*

Briefly, cells harbouring the corresponding Ll.LtrB pSEVA derivative plasmid were grown in LB supplemented with the corresponding antibiotics. When an OD of 0.2 was reached, the right inducers were added to the medium and cells were incubated at 30°C for different periods (from 30 min to 4h depending on the expression system). When IPTG was used, cells were washed and recuperated in fresh media after the induction period. Finally, serial dilutions were plated to assess viability and intron insertion efficiency on selective media when possible.

### Ll.LtrB insertion assay in *P. putida*

The same protocol described above for *E. coli* was used with *P. putida* strains with the only difference that the induction time was 2h or 4h (depending on expression system) and no recovery was performed after induction. Also, as *pyrF* gene was the target of Ll.LtrB insertion, cells were plated on M9 minimal media supplemented with only 20 μg/mL uracil (Ura) to assess viability or uracil and 250 μg/mL 5FOA (5-Fluoroorotic acid) to counter select *pyrF*-disruption mutants and make easier the identification of insertion events.

### CRISPR/Cas9 counterselection assay in *P. putida* KT2440 and KT2440 Δ*recA*

When CRISPR/Cas9 counterselection was to be applied, the protocol was adapted to simplify the process. Cells harbouring both pSEVA421-Cas9tr and pSEVA6511-GIIi derivative were grown overnight at 30°C. Next day, 1 mM cyclohexanone was added to the culture and cells were induced for 4h at 30°C. After this incubation, 1mL of cells were plated on M9 minimal media supplemented with uracil and 5FOA to assess the native efficiency of insertion in this condition. Later, cells were made electrocompetent and 100 ng of pSEVA231-CRISPR or pSEVA231-C-spacer (pyrF1, 37s or 94a depending on the experiment) were electroporated. Finally, cells were recovered in LB/Sm for 2h at 30°C, period after which serial dilutions were plated on LB/Sm (to assess viability) and LB/Sm/Km (to assess counterselection efficiency).

### Analysis of Ll.LtrB insertion by colony PCR

Ll.LtrB integrations were studied by colony PCR to check the presence or absence of the intron in the correct loci. Two possible reactions were used: In one, primers flanking the insertion site to amplify the whole intron were used. The product of this PCR would be composed of the intron sequence and the amplified flanking regions. In the second, one primer annealed in the target locus and the other inside the intron sequence, consequently a PCR product was only obtained when Ll.LtrB intron was present. In the case of barcode delivery, pool PCR reactions with a total of 4 colonies per reaction were set first. PCRs were analyzed by electrophoresis on agarose gel and 1xTAE (Tris-Acetate-EDTA). EZ Load 500bp Molecular Ruler (Brio-Rad) was the DNA ladder in all gels.

## Associated content

### Supporting information

**Supplementary Table S1:** List of oligonucleotides used in this study

**Supplementary Table S2:** List of plasmids used in this work

**Supplementary Table S3:** Insertion frequencies of Ll.LtrB::Lux1 and Ll.LtrB::Lux4 in *P. putida* KT2440 WT and Δ*recA* with no CRISPR/Cas9-mediated counterselection.

**Supplementary Table S4:** Insertion frequency of Ll.LtrB::LuxN intron in *P. putida* KT2440 WT with 5FOA CRISPR/Cas9-mediated counterselection.

**Supplementary Table S5:** Insertion frequency of Ll.LtrB::LuxN intron in *P. putida* KT2440 Δ*recA* with 5FOA CRISPR/Cas9-mediated counterselection.

**Supplementary Figure S1:** pSEVA plasmids for the expression of Ll.LtrB intron in a wide range of Gram-negative bacteria.

**Supplementary Figure S2:** Assessing size-restriction of intron-mediated delivery using *luxC* fragments as cargo and 5-FOA counterselection in log-phase induced cells.

**Supplementary Figure S3:** Barcode generation with a PCR using 3’-overlapping 119-mer oligonucleotides.

**Supplementary Figure S4:** Application of GIIi as a barcode delivery system.

**Supplementary Figure S5:** Design and test of Locus 1 and 2 spacers for CRISPR/Cas9-mediated counterselection of Ll.LtrB::B3 group II intron.

## AUTHOR INFORMATION

### Authors

Elena Velazquez— Systems and Synthetic Biology Department, Centro Nacional de Biotecnología (CNB-CSIC), Campus de Cantoblanco, Madrid 28049, Spain. orcid.org/0000-0002-0133-0461

Yamal Al-Ramahi— Systems and Synthetic Biology Department, Centro Nacional de Biotecnología (CNB-CSIC), Campus de Cantoblanco, Madrid 28049, Spain.

Jonathan Tellechea-Luzardo – Interdisciplinary Computing and Complex Biosystems (ICOS). Research Group, Newcastle University, Newcastle Upon Tyne NE4 5TG, U.K. orcid.org/0000-0003-2198-4558

Natalio Krasnogor – Interdisciplinary Computing and Complex Biosystems (ICOS) Research Group, Newcastle University, Newcastle Upon Tyne NE4 5TG, U.K.; orcid.org/0000-0002-2651-4320

### Author Contributions

EV, VdL and NK planned the experiments; EV and JT did the practical work. All Authors analyzed and discussed the data and contributed to the writing of the article.

### Notes

The authors declare no competing financial interest

