## Supplementary Information for "Targetron-assisted delivery of exogenous DNA sequences into *Pseudomonas putida* through CRISPR-aided counterselection"

by

Elena Velázquez[1], Yamal Al-Ramahi[1], Jonathan Tellechea[2], Natalio Krasnogor[2] and Víctor de Lorenzo[1]*

*[1] Systems and Synthetic Biology Department, Centro Nacional de Biotecnología (CNB-CSIC), Campus de Cantoblanco, Madrid 28049, Spain. [2]Interdisciplinary Computing and Complex Biosystems (ICOS) Research Group, Newcastle University, Newcastle Upon Tyne NE4 5TG, U.K.*

**Supplementary Table S1.** List of oligonucleotides used in this study

| Name | Sequence (5' → 3')* | Tm (°C) | Use |
|---|---|---|---|
| pS1 | AGG GCG GCG GAT TTG TCC | 71.7 | Sequencing of pSEVA from Terminator T1 |
| pS2 | GCG GCA ACC GAG CGT TC | 70.6 | Sequencing of pSEVA from Terminator T0 |
| T7p | TAA TAC GAC TCA CTA TAG GG | 50.9 | Sequencing from T7 promoter region |
| pLacZ_F | CTC GTT GCT GCA TAA ACC GAC | 66.6 | PCR to verify Ll.LtrB insertion in locus 1063a of *LacZ* gene |
| pLacZ_R | GAT GGA CCA TTT CGG CAC AGC | 70.9 | |
| pGIIintron_fwd | TTA TTA TA**T TAA TTA A**CG CGA AAT TAA TAC GAC TCA C | 66.1 | Construction of pSEVA421-GIIi |
| pGIIintron_rev | TTA TTA TA**A CTA GT**G GTG CGG ACT GTT GTA ACT C | 68.4 | |
| pSEVA427-out-SpeI | TGC GTT CGG TCA AGG TTC | 65.1 | Sequencing pSEVA421-GIIi |
| pAR1219_fwd | TTA TTA TA**T TAA TTA A**CA GAT CCC GGA CAC CAT CGA ATG GCG CAA AAC C | 81.4 | Construction of pSEVA131-T7RNAP |
| pAR1219_rev | TTA TTA TA**A CTA GT**G GCG TTA CGC GAA CGC GAA GTC CGA CTC TAA GAT G | 81.4 | |
| EBS universal | CGA AAT TAG AAA CTT GCG TTC AGT AAA C | 66.2 | Universal primer for Ll.LtrB intron retargeting |
| pyrF_165a-EBS1d | CAG AT**T GTA CA**A ATG TGG TGA TAA CAG ATA AGT CGA AGC CCT TAA CTT ACC TTT CTT TGT | 80.3 | Ll.LtrB retargeting for insertion in locus 165a (antisense orientation) of *P. putida pyrF* gene. In combination with primer EBS universal |
| pyrF_165a-IBS | AAA A**AA GCT T**AT AAT TAT CCT TAC ACT TCG AAG CCG TGC GCC CAG ATA GGG TG | 82.9 | |
| pyrF_165a-EBS2 | TGA ACG CAA GTT TCT AAT TTC GGT TAA GTG TCG ATA GAG GAA AGT GTC T | 79.6 | |
| pyrF_F | ACT TGC CAA GAG ACC CTG | 60.5 | |

| | | | |
|---|---|---|---|
| pyrF_R | TCA CGC CAA TCA GCA ACG | 67.4 | Confirmation of insertion of Ll.LtrB intron in locus 165a |
| pRetarget-fwd | AGA GTC GAC CTG CAG GCA TGC **AAG CTT** ATA ATT ATC CTT A | 77.6 | Construction of retargeted pSEVA2311-GIIi and pSEVA6511-GIIi derivatives |
| pRetarget-rev | GTT CTC CTA CAG AT**T GTA CA**A ATG TGG TGA TAA CAG ATA | 73.0 | |
| Barcode_3_F | CGT CCA GAT ATT TAT TAC GTG GCG **ACG CGT** TGG ACA TAC ATA GTA TAC TCT GGT GTT GAA GTT TCA AGG TTT TAT CCG TAG GTT CAA CTG CGT TGG GAG TGG GCA AGG GGG AAT GGG CA | 94.4 | Construction of pSEVA6511-GIIi(B3) |
| Barcode_3_R | TTT CGC TAT CAT TGC CAT TTC CCA **ACG CGT** CCA AAG GCA AAC TTG TCA AGT CTA TGG ATA ACC TAC TAA CCC CAT TGT TTT TAT ATC TAT GCC CAT TCC CCC TTG CCC ACT CCC AAC GC | 94.2 | |
| spacer 94a-F | [Phos] AAA CTG TCG CAG GAG CCG GTG AGG TCT GTC TCA TG | 83.5 | Construction of pSEVA231-C-94a |
| spacer 94a-R | [Phos] AAA ACA TGA GAC AGA CCT CAC CGG CTC CTG CGA CA | 83.6 | |
| spacer 37s-F | [Phos] AAA CCA TGG ATT GCT GGT CCC TAG GGA GCA GGA AG | 81.4 | Construction of pSEVA231-C-37s |
| Spacer 37s-R | [Phos] AAA ACT TCC TGC TCC CTA GGG ACC AGC AAT CCA TG | 80.1 | |
| 37s-IBS | AAA A**AA GCT T**AT AAT TAT CCT TAT AGG GCG CAG GAG TGC GCC CAG ATA GGG TG | 83.1 | Ll.LtrB retargeting for insertion in locus 37s (sense orientation) between *PP5408* and *glmS* genes in *P. putida.* In combination with primer EBS universal |
| 37s-EBS1d | CAG AT**T GTA CA**A ATG TGG TGA TAA CAG ATA AGT CGC AGG AAA TAA CTT ACC TTT CTT TGT | 79.9 | |
| 37s-EBS2 | TGA ACG CAA GTT TCT AAT TTC GGT TCC CTA TCG ATA GAG GAA AGT GTC T | 80.7 | |
| 94a-IBS | AAA A**AA GCT T**AT AAT TAT CCT TAG GTG ACG TCT GTG TGC GCC CAG ATA GGG TG | 82.3 | Ll.LtrB retargeting for insertion in locus 94a (antisense orientation) between *PP5408* and *glmS* genes in *P. putida.* In |
| 94a-EBS1d | CAG AT**T GTA CA**A ATG TGG TGA TAA CAG ATA AGT CGT CTG TCT TAA CTT ACC TTT CTT TGT | 78.8 | |

| | | | |
|---|---|---|---|
| 94a-EBS2 | TGA ACG CAA GTT TCT AAT TTC GAT TTC ACC TCG ATA GAG GAA AGT GTC T | 80.6 | combination with primer EBS universal |
| pbarcodes universal | TGG ACA TAC ATA GTA TAC TCT GGT G | 58.6 | Sequencing of Barcodes and PCR for confirmation of intron insertion |
| pbarcodes GIIi reverse | ACA CAA TAA CTG TAC CCC TTT GCC | 65.7 | |
| 649-Tn7-F | CGA TTC ATC AGG TTG GAT TCG | 66.4 | PCR confirmation of Ll.LtrB insertion in *PP5408-glms* intronic region |
| 418-Tn7-R | AAT CTG GCC AAG TCG GTG AC | 66.3 | |
| pLux_fwd | TTA TTA T**AC GCG** TAT GAC TAA AAA AAT TTC ATT CAT TAT TAA CGG | 71.7 | Amplification of *LuxC* gene to produce cargos of different sizes |
| pLux1_rev | TTA TTA T**AC GCG** TAT TAC AAT CAA TAA TGT TTT TTA CAT GAG AGT C | 71.1 | |
| pLux2_rev | TTA TTA T**AC GCG** TCT CTA GCT TAG CCA TTT CTT CTG | 71.5 | |
| pLux3_rev | TTA TTA T**AC GCG** TCA GAT GTA CAG ATT TAC CTT TC | 68.8 | |
| pLux4_rev | TTA TTA T**AC GCG** TCG GAT GAT TAG GGT CTA C | 69.8 | |
| pLux5_rev | TTA TTA T**AC GCG** TAT CAG CAT AAG ATG GCG | 70.5 | |
| pLux6_rev | TTA TTA T**AC GCG** TAT GAT TTC CCA TGT AAT ATA TGT TTT G | 70.4 | |
| pLux7_rev | TTA TTA T**AC GCG** TAT GAT TTC CCA TGT AAT ATA TGT TTT G | 68.2 | |
| pgII_cargo_fwd | TAG TAG TCT GAG AAG GGT AAC G | 52.8 | Sequencing Lux inserts in Ll.LtrB intron. Confirmation of Ll.LtrB insertion in combination with primers pyrF_F and pyrF_R, respectively. |
| pGIIi_cargo_rev | GTA TAC GGC TCT GTT ATT GTT C | 51.0 | |

*Bold letters correspond to restriction enzyme sites

[Phos]: Phosphorothioate group

**Supplementary Table S2**. List of plasmids used in this work

| Plasmid | Description | Reference |
|---|---|---|
| **pAR1219** | Expression plasmid carrying T7 RNA polymerase gene (bacteriophage gene 1) under control of *lacUV5* promoter; *oriV*(pMB1); Ap$^R$ | Merck (Sigma-Aldrich) |
| **pACD4K-C** | Expression plasmid carrying Ll.LtrB intron (bearing Km$^R$ Retrotransposition-Activated Marker, RAM) under control of T7 promoter and retargeted to insert into locus 1063 of *E. coli LacZ* gene in the antisense orientiation; *oriV*(p15A); Cm$^R$ | Merck (Sigma-Aldrich) |
| **pSEVA231-CRISPR** | pSEVA231 derivative bearing CRISPR array; oriV (pBBR1); Km$^R$ | [1] |
| **pSEVA421-Cas9tr** | pSEVA421 derivative bearing the cas9 gene and tracrRNA; oriV (RK2); Sm$^R$/Sp$^R$ | [1] |
| **pSEVA131** | Standard SEVA expression vector; *oriV*(pBBR1); Ap$^R$ | [2] |
| **pSEVA427** | Standard SEVA expression vector; *oriV*(pBBR1); *gfp* cargo; Sm$^R$/Sp$^R$ | [2] |
| **pSEVA131-T7RNAP** | pSEVA131 derivative with T7 RNA polymerase gene expression under *lacUV5* promoter control | This work |
| **pSEVA421-GIIi(Km)** | pSEVA427 derivative expressing Ll.LtrB group II intron under T7 promoter control. Ll.LtrB bearing the Km$^R$ RAM (Retrotransposition-Activated selectable Marker) and retargeted to insert into 1063 locus of *E. coli LacZ* gene in antisense orientation. | This work |
| **pSEVA421-GIIi-pyrF** | pSEVA421-GIIi(Km) derivative with empty Ll.LtrB intron and retargeted to insert into 165 locus of *P. putida pyrF* gene in antisense orientation. | This work |
| **pSEVA421-GIIi(Km)-pyrF** | pSEVA421-GIIi(Km) derivative with Ll.LtrB intron bearing Km$^R$ RAM and retargeted to insert into 165 locus of *P. putida pyrF* gene in antisense orientation. | This work |
| **pSEVA2311** | Standard SEVA expression vector; *oriV*(pBBR1); *ChnR*-$P_{ChnB}$, cyclohexanone-responsive expression plasmid; Km$^R$ | [3,4] |
| **pSEVA2311-GIIi(Km)** | pSEVA2311 derivative with Ll.LtrB intron under *ChnR*-$P_{ChnB}$ promoter. Ll.LtrB intron bearing Km$^R$ RAM and retargeted to insert into 1063 locus of *E. coli LacZ* gene in antisense orientation. | This work |
| **pSEVA2311-GIIi-pyrF** | pSEVA2311-GIIi(Km) derivative with empty (no RAM) Ll.LtrB intron retargeted to insert into 165 locus of *P. putida pyrF* gene in antisense orientation. | This work |
| **pSEVA651** | Standard SEVA expression vector; *oriV*(RSF1010); Gm$^R$ | [2] |

| | | |
|---|---|---|
| **pSEVA6511-GIIi(Km)** | pSEVA651 derivative with *ChnR*-$P_{ChnB}$ promoter and Ll.LtrB intron from pSEVA2311-GIIi(Km) as a PacI/SpeI insert. | This work |
| **pSEVA6511-GIIi** | pSEVA6511-GIIi(Km) derivative with no $Km^R$ RAM (empty Ll.LtrB) | This work |
| **pSEVA6511-GIIi-pyrF** | pSEVA6511-GIIi derivative retargeted to insert into 165 locus of *P putida pyrF* gene. | This work |
| **pSEVA231-C-pyrF1** | pSEVA231-CRISPR derivative with pyrF spacer cloned into BsaI restriction sites. | [1] |
| **pSEVA256** | Standard SEVA expression vector; *oriV*(RSF1010); *luxCDABE* as cargo; $Km^R$ | [2] |
| **pSEVA6511-GIIi(Lux1)-pyrF** | pSEVA6511-GIIi-pyrF derivative with Lux1 (150 bp) insert in sense orientation | This work |
| **pSEVA6511-GIIi(Lux2)-pyrF** | pSEVA6511-GIIi-pyrF derivative with Lux2 (300 bp) insert in sense orientation | This work |
| **pSEVA6511-GIIi(Lux3)-pyrF** | pSEVA6511-GIIi-pyrF derivative with Lux3 (450 bp) insert in sense orientation | This work |
| **pSEVA6511-GIIi(Lux4)-pyrF** | pSEVA6511-GIIi-pyrF derivative with Lux4 (600 bp) insert in sense orientation | This work |
| **pSEVA6511-GIIi(Lux5)-pyrF** | pSEVA6511-GIIi-pyrF derivative with Lux5 (750 bp) insert in sense orientation | This work |
| **pSEVA6511-GIIi(Lux6)-pyrF** | pSEVA6511-GIIi-pyrF derivative with Lux6 (900 bp) insert in sense orientation | This work |
| **pSEVA6511-GIIi(Lux7)-pyrF** | pSEVA6511-GIIi-pyrF derivative with Lux7 (1050 bp) insert in sense orientation | This work |
| **pSEVA231-C-37s** | pSEVA231-CRISPR derivative with 37s spacer cloned into BsaI restriction sites | This work |
| **pSEVA231-C-94a** | pSEVA231-CRISPR derivative with 94a spacer cloned into BsaI restriction sites | This work |
| **pSEVA6511-GIIi(B3)-37s** | pSEVA6511-GIIi derivative retargeted to insert into locus 37 (in sense orientation) between *PP5408* and *glmS* genes in *P. putida* and bearing barcode 3 as a MluI insert. | This work |
| **pSEVA6511-GIIi(B3)-94a** | pSEVA6511-GIIi derivative retargeted to insert into locus 94 (in antisense orientation) between *PP5408* and *glmS* genes in *P. putida* and bearing barcode 3 as a MluI insert. | This work |

**Supplementary Table S3.** Insertion frequencies of Ll.LtrB::Lux1 and Ll.LtrB::Lux4 in *P. putida* KT2440 WT and Δ*recA* with no CRISPR/Cas9-mediated counterselection.

| | From Ura (+ intron/total re-streaks) | From FOA (+ intron/total re-streaks) | |
|---|---|---|---|
| **Lux1** | 1/105 (0.95%) | 17/108 (16%) | wt |
| **Lux4** | 0/104 | 2/50 (4%) | |
| **Lux1** | 0/101 | 10/106 (9.4%) | ΔrecA |
| **Lux4** | 0/102 | 0/61 | |

**Supplementary Table S4.** Insertion frequency of Ll.LtrB::LuxN intron in *P. putida* KT2440 WT with 5FOA CRISPR/Cas9-mediated counterselection.

| | | Replicate 1 | Replicate 2 | Replicate 3 | Total |
|---|---|---|---|---|---|
| **5FOA** | **LuxØ** | 0/58 | 1/53 | 0/51 | 1/162 |
| | **Lux1** | 0/60 | 0/56 | 0/56 | 0/172 |
| | **Lux2** | 0/60 | 0/31 | 0/51 | 0/142 |
| | **Lux3** | 0/59 | 0/56 | 0/55 | 0/170 |
| | **Lux4** | 0/60 | 0/49 | 0/54 | 0/163 |
| **231-CRISPR** | **LuxØ** | 0/10 | 2/48 | 4/46 | 6/104 |
| | **Lux1** | 0/56 | 0/2 | 0/104 | 0/162 |
| | **Lux2** | 0/42 | 0/53 | 0/107 | 0/202 |
| | **Lux3** | 0/56 | 0/56 | 0/106 | 0/2018 |
| | **Lux4** | 0/48 | 0/52 | 0/104 | 0/204 |
| **231-C-pyrF1** | **LuxØ** | 13/18 | 7/56 | 17/40 | 37/114 |
| | **Lux1** | 1/53 | 0/24 | 0/10 | 1/87 |
| | **Lux2** | 0/56 | 0/51 | - | 0/107 |
| | **Lux3** | 0/57 | 0/50 | 0/96 | 0/203 |
| | **Lux4** | 0/56 | 0/48 | 0/87 | 0/191 |

*Number of positive intron-insertion colonies confirmed through PCR divided by total of screened colonies.

**Supplementary Table S5.** Insertion frequency of Ll.LtrB::LuxN intron in *P. putida* KT2440 Δ*recA* with 5FOA CRISPR/Cas9-mediated counterselection.

| | | Replicate 1 | Replicate 2 | Replicate 3 | Total |
|---|---|---|---|---|---|
| 5FOA | LuxØ | 0/35 | 3/51 | 5/28 | 8/114 |
| | Lux1 | 0/50 | 0/56 | 0/56 | 0/162 |
| | Lux2 | 0/38 | 0/50 | 0/49 | 0/137 |
| | Lux3 | 0/40 | 0/54 | - | 0/94 |
| | Lux4 | 0/56 | 0/51 | - | 0/107 |
| 231-CRISPR | LuxØ | 1/51 | 3/30 | 1/49 | 5/130 |
| | Lux1 | 1/55 | 1/34 | 0/50 | 2/139 |
| | Lux2 | 0/56 | 0/35 | 0/53 | 0/144 |
| | Lux3 | 0/54 | 0/56 | - | 0/110 |
| | Lux4 | 0/53 | 0/39 | - | 0/92 |
| 231-C-pyrF1 | LuxØ | 1/40 | 7/40 | 4/40 | 11/120 |
| | Lux1 | 2/48 | 1/40 | 10/40 | 13/128 |
| | Lux2 | 7/15 | 1/40 | 2/3 | 10/58 |
| | Lux3 | 3/23 | 3/10 | - | 6/33 |
| | Lux4 | 2/8 | 1/1 | - | 3/9 |

*Number of positive intron-insertion colonies confirmed through PCR divided by total of screened colonies.

**Supplementary Fig. S1.** pSEVA plasmids for expression of Ll.LtrB intron in a wide range of Gram-negative bacteria.

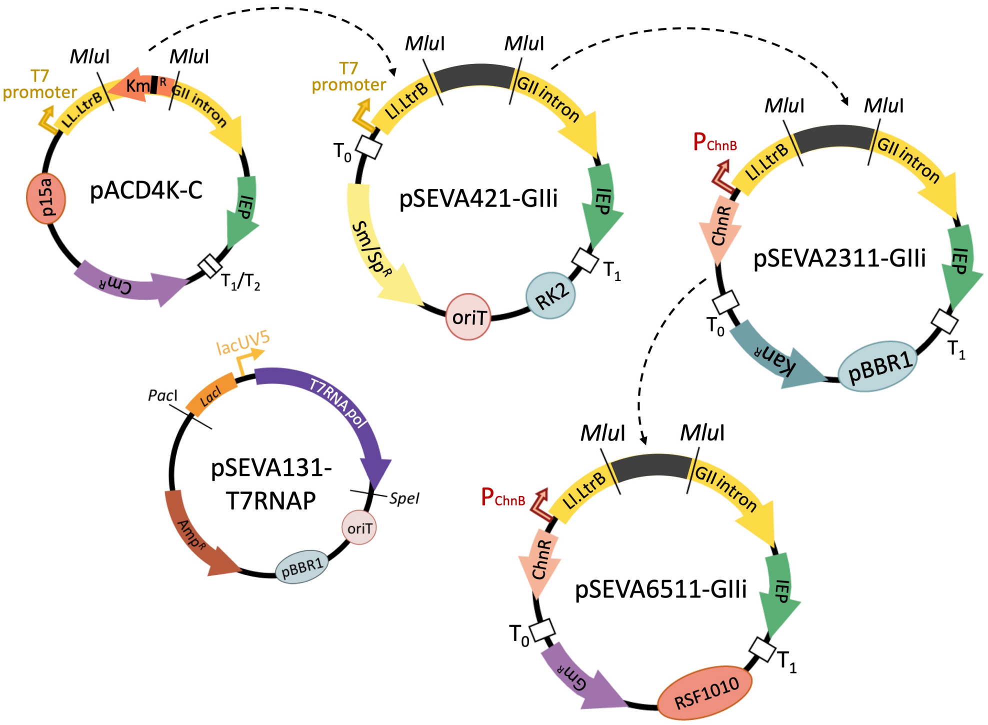


General structure and features of the main vectors used in this study are shown starting with the original plasmid (pACD4K-C, Sigma-Aldrich) used for the amplification and subcloning of Ll.LtrB.

**Supplementary Fig. S2.** Assessing intron-mediated delivery of DNA fragments.

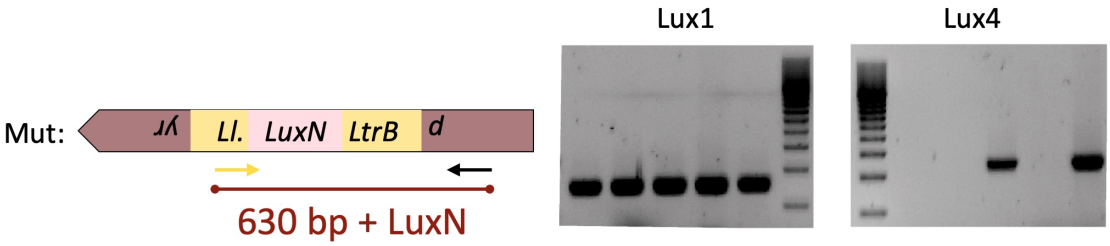


The figure to the left sketches the genomic segment (*pyrF* gene) where *lux* fragments were loaded as a cargo, next to size-restriction analysis of the PCR of some colonies after log-phase induced cells followed of 5-FOA counterselection.

**Supplementary Fig. S3.** Barcode generation with a PCR using 3'-overlapping 119-mer oligonucleotides.

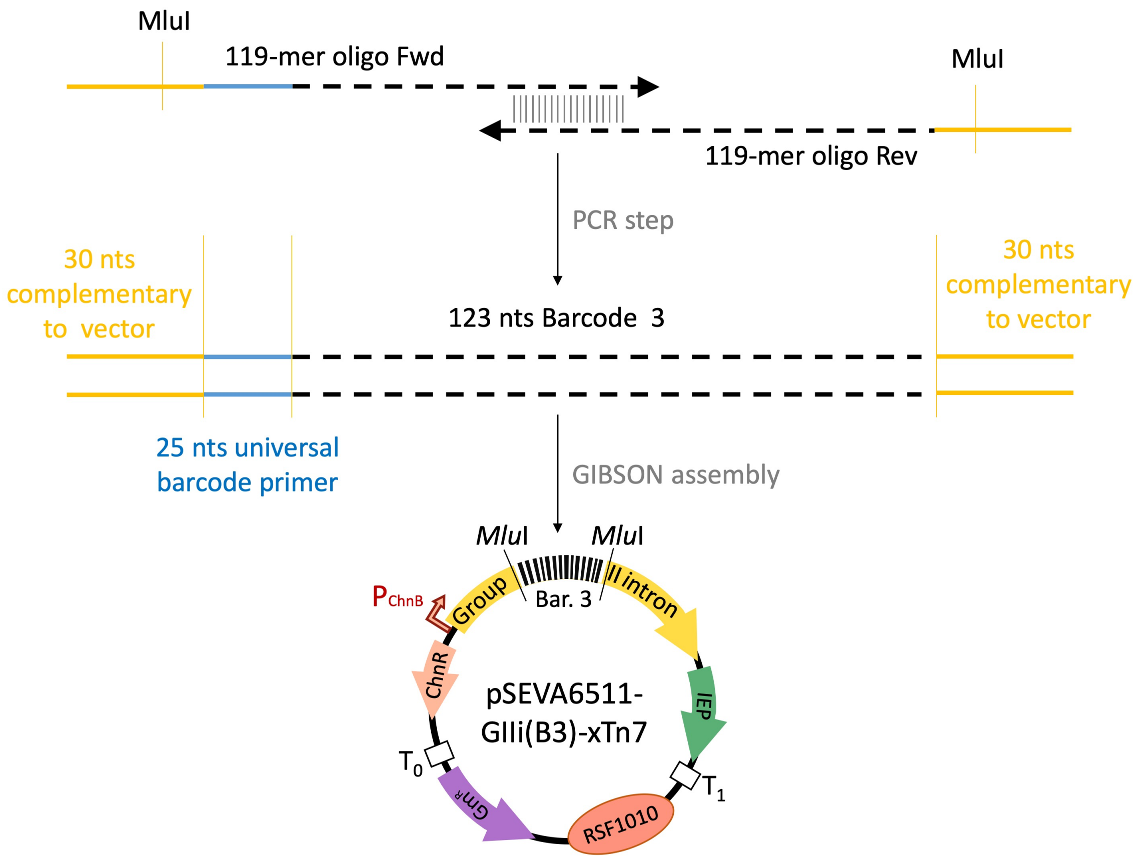


Once generated, the PCR fragment (208 bp) was directly assembled into linearized-pSEVA6511-GIIi and then this plasmid was retargeted either to the 37s locus (pSEVA6511-GIIi(B3)-37s) or to the 94a locus (pSEVA6511-GIIi(B3)-94a)

**Supplementary Fig. S4.** Application of GIIi as a barcode delivery system.

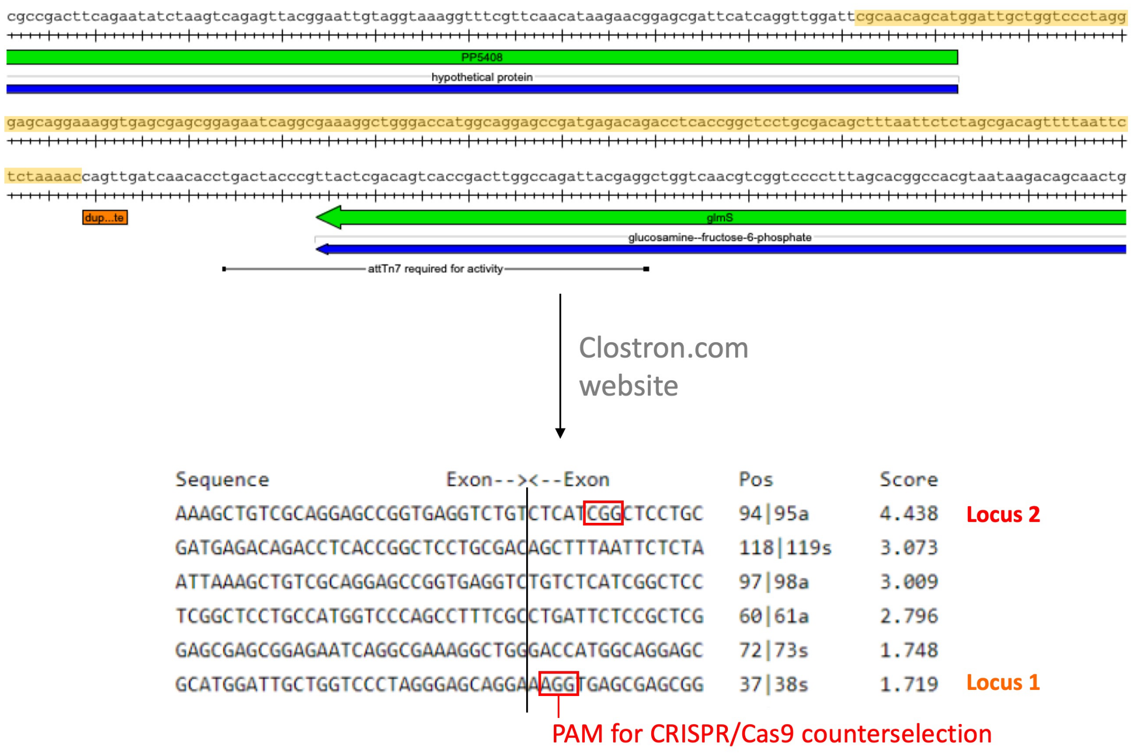


Selection of the insertion loci to be tested for the retargeting of Ll.LtrB to the close vicinity of Tn7-insertion site. Important sequences for insertion of Tn7 transposon are featured in the figure (orange box and black line). The region used in the clostron website to look for suitable insertion loci is highlighted in yellow. The retrieved list with posible retargeting loci is shown along with the two selected loci. Note the red boxes emphasizing the PAM sequences necessary for CRISPR/Cas9-mediate counterselection and used as the starting point for the design of corresponding spacers.

**Supplementary Fig S5.** Design and test of Locus 1 and 2 spacers for CRISPR/Cas9-mediated counterselection of Ll.LtrB::B3 group II intron.

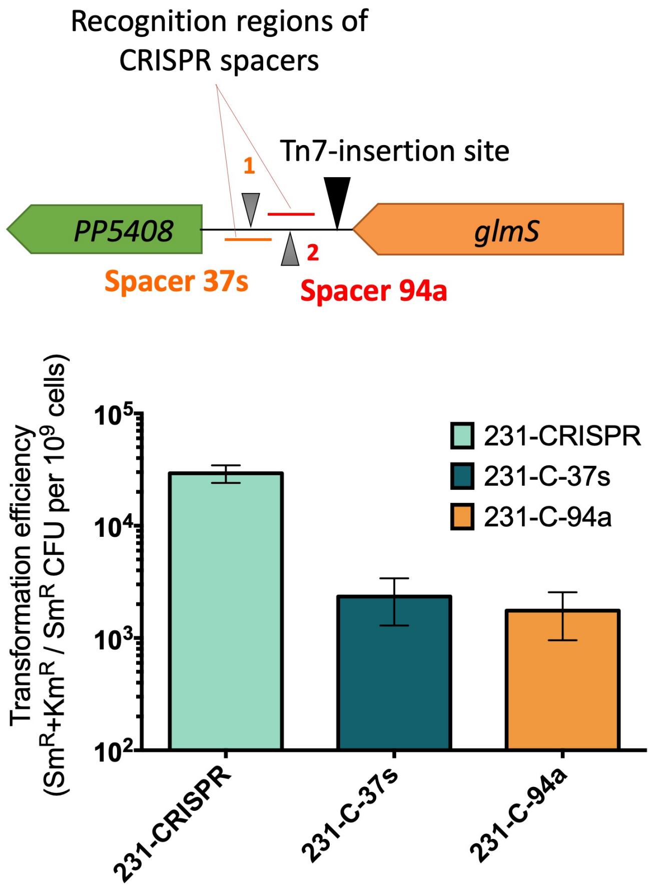


(Top) Recognition región of spacer 37s (orange) and 94a (red). Locus 1 and 2 are indicated with grey trangles. Tn7-insertion site is shown with a black triangle. (Bottom) Efficiency of cleavage of spacer 37s and 94a in comparison to control spacer present in pSEVA231-CRISPR.
